# Akt Drives TGF-β-induced Over-secretion of DKK1 and Impairment of Cystic Fibrosis Airway Epithelium Polarity

**DOI:** 10.1101/2023.05.22.541703

**Authors:** Tahir Idris, Michaël Bachmann, Marc Bacchetta, Wehrle-Haller Bernhard, Marc Chanson, Mehdi Badaoui

## Abstract

Epithelial polarity is fundamental in maintaining barrier integrity and tissue protection. In cystic fibrosis (CF), apicobasal polarity of the airway epithelium is lost, resulting in increased apical fibronectin deposition and enhanced susceptibility to bacterial infections. Here we show that *CFTR* mutation in primary human airway epithelial cells (HAECs) and *CFTR* knockdown in a HAEC line promote the overexpression and over-secretion of TGF-β1 and DKK1 when cultured at air-liquid interface (ALI). These dynamic changes result in hyperactivation of the TGF-β pathway and inhibition of the Wnt pathway through degradation of β-catenin leading to imbalanced proliferation and polarization. The abnormal interplay between TGF-β and Wnt signaling pathways is further enhanced by aberrant Akt signaling. Pharmacological manipulation of TGF-β, Wnt, and Akt pathways restored polarization of the CF epithelium. Our data shed new insights into the signaling pathways that fine-tune apicobasal polarization and also highlight new therapeutic strategies in preventing infections in CF HAECs.

## INTRODUCTION

Proper polarization and differentiation are necessary to shape an epithelium and to maintain its structural integrity. The airway epithelium is frequently exposed to injury-causing agents and its regeneration is crucial in maintaining barrier function. Indeed, defects in the polarization and differentiation processes can alter the regeneration of an epithelium resulting in a lack of integrity and barrier function which can be manifested in several disorders ^1^.

Cystic fibrosis (CF) is an autosomal, recessive genetic disorder caused by CF transmembrane conductance regulator (CFTR) mutant proteins that result in an impairment of chloride and bicarbonate ion transport ^2, 3^. This causes several pulmonary manifestations including chronic bacterial infections, exacerbated inflammation and mucus plugging of the trachea-bronchial airway tract, leading to bronchiectasis and eventually respiratory failure ^2, 3^.

CF airway disease can be attributed to a loss of cell polarity and loss of barrier integrity thereby predisposing the epithelium to increased bacterial infections ^4^. Airway epithelial apicobasal polarity is dependent on a well-organized cytoskeletal, junctional network and proper establishment of extracellular matrix (ECM) proteins. In CF, an alteration and disorganization of cytoskeletal and junctional complexes have been reported in multiple studies ^5–9^. Furthermore, we previously reported that CF airway epithelia display an overexpression and ectopic apical localization of the ECM protein fibronectin and its receptor β1-integrin, leading to increased *Pseudomonas aeruginosa* (*Pa*) adhesion to the CF airways ^10, 11^. Additionally, CF airways have been shown to manifest a partial epithelial-to-mesenchymal transition (EMT) phenotype, suggesting an abnormal differentiation of the epithelium ^12, 13^. Altogether, these reports hint towards a defective polarization process of the CF epithelia. However, the mechanistic details await further exploration.

Several signaling pathways are crucial in maintaining proper epithelial polarization and differentiation in the airways. The Wingless-related integration site (Wnt), Transforming Growth Factor β (TGF-β) and Akt signaling pathways and their interplay have been well described to play important roles in both these processes in multiple contexts ^14–16^. Inhibition of the canonical Wnt pathway results in the degradation of β-catenin by the so-called destruction complex that includes glycogen synthase kinase (GSK)-3β among other proteins ^17^. This inhibited state is promoted by the actions of dickkopf-related protein 1 (DKK1), a secreted protein that binds to the low-density lipoprotein receptor-related protein (LRP) 6 co-receptor. Activation of the pathway by Wnt ligands inhibits this complex and allows β-catenin to activate its target genes ^17^. Wnt signaling is known to be involved in the regeneration process of the airway epithelium after injury. Indeed, the regeneration process is dependent on proliferation, polarization, and terminal differentiation of the different cell types of the airways. Wnt signaling has been shown to promote proliferation and renewal of the airway basal stem cells thus inhibiting differentiation into ciliated cell lineage ^18^. In another context, DKK1 was shown to localize to adhesion complexes and represses cell polarization and integrity of cell-cell adhesion in a zebrafish model ^19^. Despite the well-described effects of the Wnt pathway on polarization in various cellular models, its role in CF airway epithelia is yet to be elucidated.

The canonical TGF-β pathway is activated by TGF-β1, TGF-β2 and TGF-β3. Activation of the pathway results in intracellular phosphorylation of Smad2 and Smad3 which, together with Smad4, translocate to the nucleus and activate TGF-β target genes ^15^. Importantly, the pathway is inhibited by Smad7, a target gene of the pathway, through a negative feedback regulation ^15^. The role of TGF-β during regeneration has also been well elucidated. For instance, TGF-β signaling can lead to remodeling of the ECM to trigger the migration process of different epithelial cell types after an injury ^20–22^. Additionally, differentiation of airway basal cells can also be triggered by Smad signaling ^23^. Importantly, TGF-β1 and TGF-β signaling are regulators of EMT and their activation can induce EMT-associated transcription factors (EMTa-TFs), including Snail and Twist1, ultimately resulting in a loss of epithelial proteins ^12, 24^. Interestingly, *TGF-β1* is a modifier gene in CF ^25^ and there is evidence of its increase in plasma and bronchoalveolar lavage fluid from individuals with CF ^26^. However, direct link between TGF-β signaling and CF disease progression is not well established.

The phosphoinositide-3 kinase/Akt pathway (PI3K/Akt) is a well-known regulator of proliferation, cell motility and wound closure during the regeneration process ^27^. Indeed, inhibiting phosphatase and tensin homolog (PTEN), an inhibitor of the pathway, led to an acceleration of wound closure in primary human airway epithelial cells (HAECs) ^28^. Conversely, inhibition of PI3K/Akt or an expression of dominant negative form of PI3K led to a decrease in cell migration after injury in the 16HBE14o− airway epithelial cell line ^29^. In the F508del homozygous CFBE41o-cell line, PI3K/Akt signaling has been linked to autophagy through the mTOR pathway ^30^, however studies on its role in CF airway biology are limited.

In CF, there is growing evidence of defects in proliferation, polarization, and differentiation processes of the epithelium ^31, 32^. Given the role of the Wnt, TGF-β and Akt signaling pathways and their interplay in regulating these processes, they appear as promising targets for investigation in CF. In this study, we report aberrant Wnt, TGF-β and Akt signaling that results in defective polarization of the CF airway epithelium. Defects or lack of CFTR led to an overexpression and over-secretion of TGF-β1 and DKK1 promoting a hyperactive TGF-β signaling and a decreased Wnt signaling, creating an imbalance between these two pathways. We also show that an aberrant Akt signaling modulates the TGF-β and Wnt pathways in CF and serves as an important mediator of both pathways. Finally, by targeting these pathways in CF HAECs with pharmacological drugs, we were able to prevent apical fibronectin deposition, an important phenotype that is symptomatic of a defect in polarity in CF cells.

## RESULTS

### Dysregulated expression of genes related to the TGF-β and Wnt signaling pathways in repairing CF HAECs

We previously performed RNA sequencing (RNA-seq) on repairing primary HAECs from CF (N=7) and non-CF (NCF, N=6) donors ^33^. We focused on the initial steps of repair by comparing conditions of early post-wounding (pW), wound closure (WC) and 2-days post wound closure (pWC) to the control conditions of each stage (Figure S1A). Gene set enrichment analyses (GSEA) of the data predicted several pathways to be significantly dysregulated. From these, three pathways, including the ‘Wnt Signaling Pathway’, Hippo Signaling Pathway,’ and ‘Basal Cell Carcinoma’ were enriched in all the phases of wound repair (Figure 1A). ‘Wnt Signaling Pathway and Pluripotency’ was enriched in the first 24 hours after wounding, ‘BMP Signaling Pathway in Eyelid Development’, was enriched during wound closure, and the TGF-β Signaling Pathway was enriched 48 hours after wound closure (Figure 1A). Within these pathways, a large number of genes with significantly higher expression in CF are shown in Figure 1A, including, interestingly, *TGF-β1* and *DKK1* in the TGF-β and Wnt signaling pathways respectively, 48 hours after wound closure (Figure 1A). To confirm these predictions, the mRNA expression of *TGF-β1* and *DKK1* was monitored by RNAscope® in fully differentiated primary CF and NCF HAECs. Both *TGF-β1* and *DKK1* were overexpressed in the CF airway epithelia (Figures 1B, C), suggesting a dysregulation of the TGF-β and Wnt signaling pathways.

**Figure 1.**
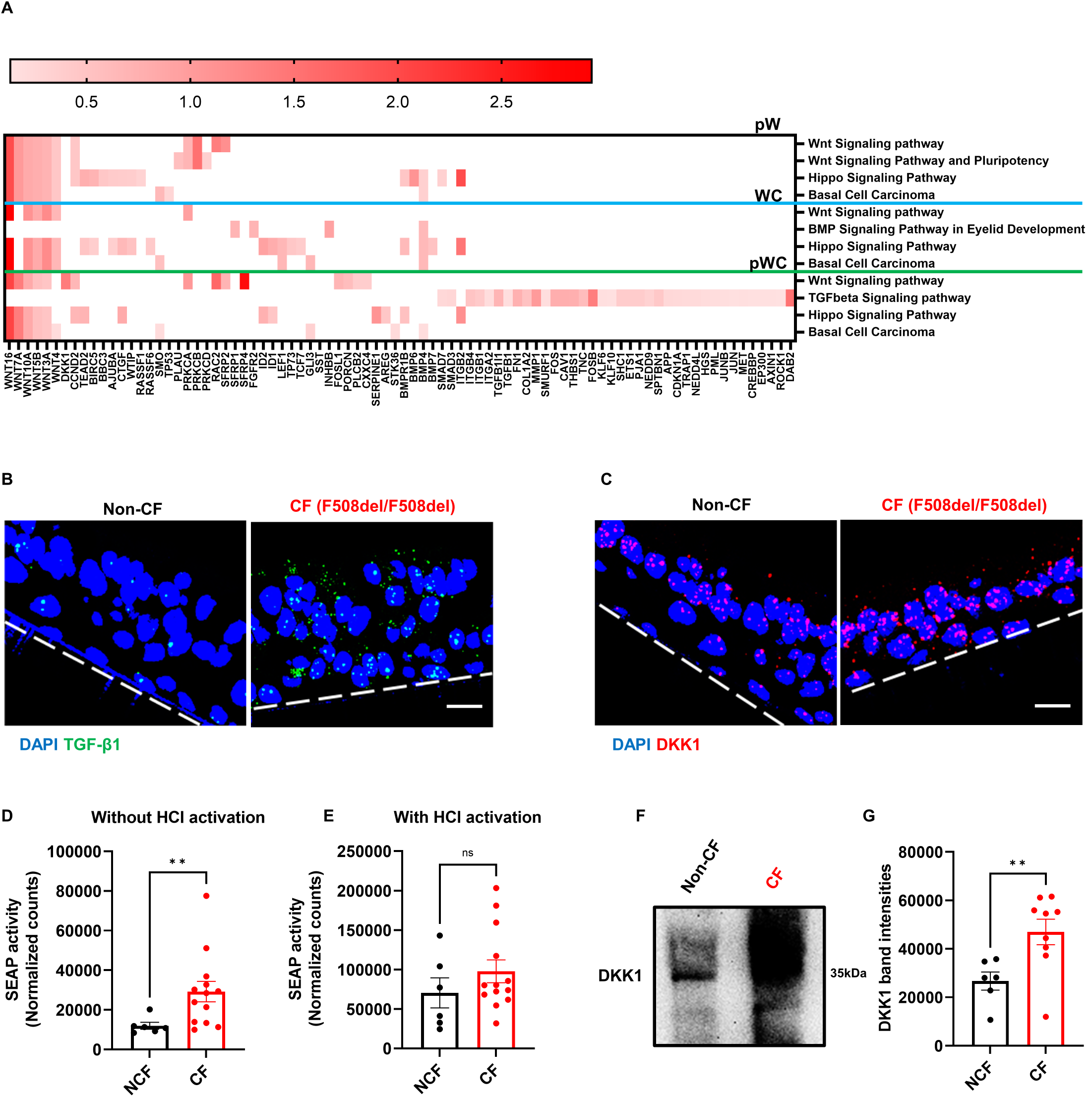
Dysregulated expression of genes related to the TGF-β and Wnt signaling pathways in repairing CF HAECs. **(A)** Heat map showing RNA-seq predictions from GSEA analysis of dysregulated genes and pathways of the repairing primary NCF (N=6) and CF (N=7) HAECs. Dysregulated pathways have a false discovery rate (FDR) <0.05 and a normalized enrichment score (NES) > 1. Significantly enriched genes are ranked by their Rank Metric Score (fold change). pW = 24 hours post wound, WC = wound closure, pWC = 48 hours post wound closure. **(B and C)** Representative RNAscope® *in situ* hybridization depicting TGF-β1 (**B**, green), DKK1 (**C**, red) mRNA expression and DAPI (blue) in fully differentiated primary NCF (N=4) and CF (N=3) HAECs. **(D and E)** SEAP activity of TGF-β reporter cell line after treatment with NCF (N=4, n=6) and CF (N=6, n=13) conditioned buffers for 24 hours without **(D)** and with **(E)** HCl activation. Mann-Whitney U test, ** p<0.01. **(F and G)** Representative western blot **(F)** and its quantification **(G)** of DKK1 secretions in primary NCF (N=4, n=6) and CF (N=6, n=9) HAECs. Mann-Whitney U test, ** p<0.01. Scale bars: 20 µm. Dashed lines in **B** and **C** depict basal side. See also Figures S1A and B.

Next, we monitored the functional consequence of the transcriptional overexpression of TGF-β1 and DKK1 in CF HAECs. To this end, as both genes are encoding for secreted proteins, we incubated differentiated primary NCF (N=4) and CF HAECs (N=6) with a physiological saline buffer for 24 hours (Figure S1B). First, we monitored TGF-β secretion and activation from NCF and CF primary HAECs after buffer incubation by using a secreted alkaline phosphatase (SEAP)-based TGF-β reporter cell line. Latent TGF-β levels were measured after hydrochloric acid (HCl) activation, while the pre-activated form was determined without HCl. We observed increased activation of TGF-β by CF HAECs (p<0.01; Figure 1D), while the total amount of secreted TGF-β was not significantly changed between NCF and CF HAECs after HCl activation (Figure 1E), demonstrating that CF HAECs have an increased ability to activate TGF-β. Next, we monitored DKK1 secretion by western blot after buffer incubation. Interestingly, quantification of DKK1 western blot revealed a significantly higher secretion in primary CF HAECs (p<0.01; Figures 1F, G), further suggesting a dysregulated Wnt signaling. These results indicate that differentiated primary CF HAECs display a hyperactive TGF-β signaling and suggest a DKK1-inhibited Wnt signaling. As these pathways are critical for tissue health and diseases ^15, 17^, we hypothesized that they may contribute to an abnormal CF airway epithelial homeostasis.

### *CFTR* knockdown leads to defective polarization and proliferation in Calu-3 cells

To investigate if the polarization process is CFTR-dependent, we compared the Calu-3 HAEC line knocked-down for *CFTR* by CRISPR-Cas9 (*CFTR* KD) to control (CTL) counterparts, grown at an air-liquid interface (ALI) as a trigger for polarization through inhibition of proliferation (Figure S1C).

We have previously shown that fully polarized CFTR KD cells cultured at an ALI displayed an overexpression and an ectopic apical appearance of fibronectin ^10, 34^. As polarization is a dynamic process, we thus performed western blotting of total fibronectin expression in both CTL and CFTR KD cells at different days of ALI culture, and this revealed an increased expression from day 1 (D1) of ALI in CFTR KD cells (p<0.05, p<0.01; Figures 2A, B). Interestingly, immunostaining revealed no apical fibronectin in both CTL and CFTR KD cells from days 0 to 2 (D0-D2) (Figure 2C). However, CFTR KD cells began to display apical fibronectin from day 3 (D3), which gradually increases until day 14 (D14). The CTL cells on the other hand did not display any ectopic appearance of fibronectin during ALI (Figure 2C). We next monitored the proliferation behavior of CTL and CFTR KD cells during ALI culture as revealed by immunostaining of Ki67, a nonhistone nuclear protein that is a biomarker for this process (Figure 2D). Quantification of the staining revealed a peak in proliferation at D3 of ALI in CTL cells that is absent in CFTR KD cells (p<0.01; Figure 2E). After D3, proliferation decreased in both CTL and CFTR KD cells. Even though the CTL cells continued proliferating until day 11 (D11), the CFTR KD cells, however, stopped proliferating at day 8 (D8) of ALI (p<0.01; Figure 2D, E). These results suggest that the lack of CFTR in a HAEC cell line leads to disruption of the proliferation/polarization balance that normally occurs during ALI.

**Figure 2.**
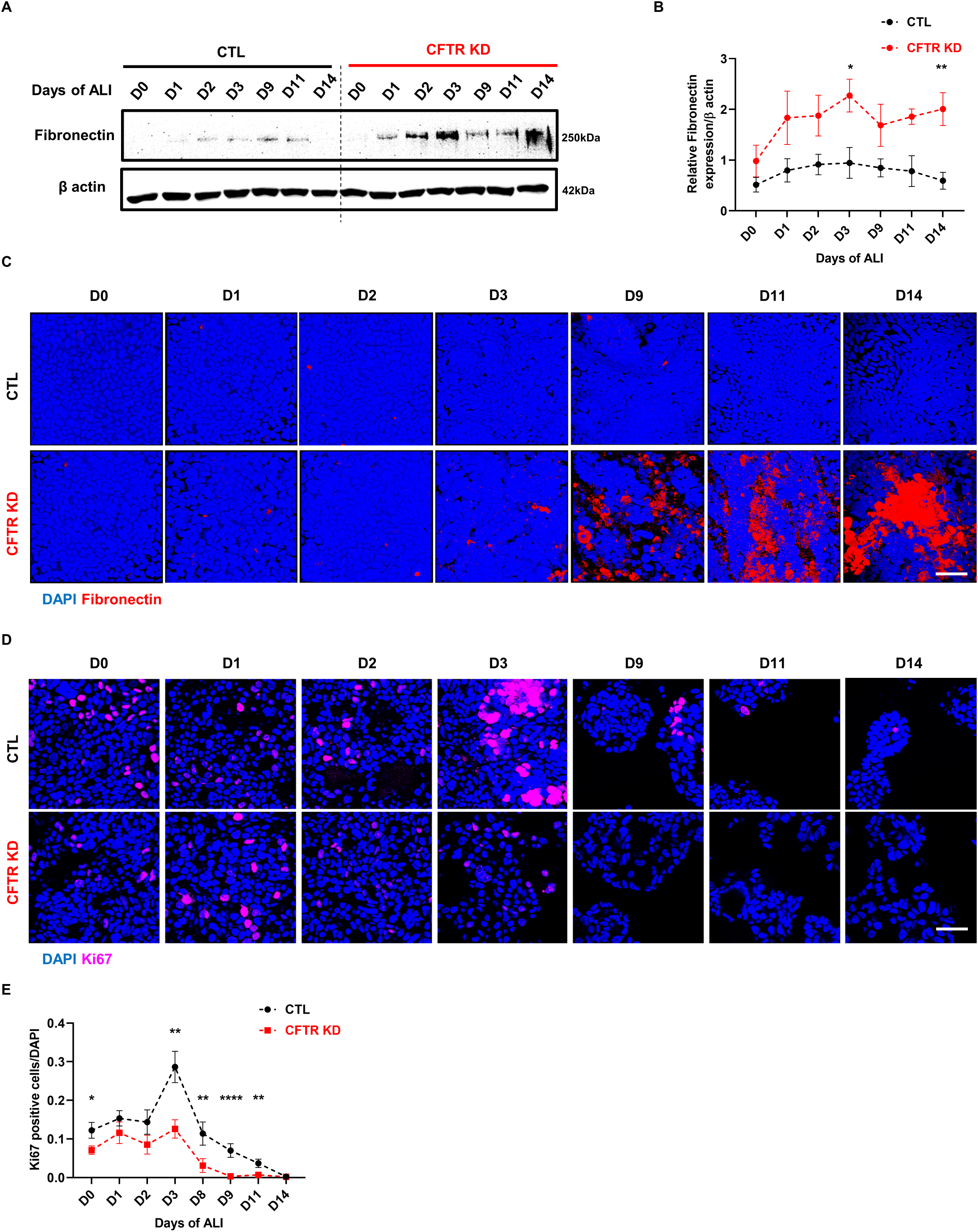
*CFTR* knockdown leads to defective polarization and proliferation in Calu-**3 cells.** **(A and B)** Representative western blot **(A)** and corresponding quantification **(B)** depicting fibronectin protein expression during ALI culture in CTL and CFTR KD cells (n=6). β-actin was used as a loading control. Mann-Whitney U test, * p<0.05, ** P<0.01. **(C)** Top view of 3D reconstructions of z-stack confocal images depicting fibronectin (red) and DAPI (blue) in CTL and CFTR KD cells during ALI (n=3). **(D and E)** Confocal images **(D)** and corresponding quantification **(E)** depicting Ki67 (magenta) and DAPI (blue) at the apical side of CTL and CFTR KD cells during ALI (n=3). Mann-Whitney U test, * p<0.05, ** p<0.01, **** p<0.0001. Scale bars: 40 µm (upper panel), 50 µm (lower panel). See also Figure S1C.

### *CFTR* knockdown leads to over-production of TGF-β1 and over-stimulation of the TGF-β pathway

There is a well-known regulatory relationship between TGF-β pathway and fibronectin expression in several cell types ^35^. *TGF-β1* gene and the TGF-β Signaling Pathway were predicted to be significantly enriched by GSEA of RNA-seq (Figure 1A), whilst an increased expression and secretion of TGF-β1 was detected in differentiated CF epithelia (Figures 1B and 1D). Therefore, we investigated if TGF-β1 production, secretion and TGF-β signaling are also dysregulated in CFTR KD cells. We first determined the level of *TGF-β1* mRNA expression by qPCR in our Calu-3 cell model, which was increased in CFTR KD cells at D14 of ALI culture (p<0.05; Figure S2A). Next, we apically incubated polarized CTL and CFTR KD cells at D14 with a physiological saline buffer for 24 hours to collect secreted TGF-β1 and treated the TGF-β reporter cell line with these buffers. Indeed, CFTR KD cells showed increased secretion (p<0.0001, Figures 3A) and increased activation of TGF-β (p<0.05; Figures 3B). We also determined TGF-β activity during ALI culture of CTL and CFTR KD cells. Interestingly, the increased response in CFTR KD cells was observed from D3 of ALI (p<0.01; Figure 3C), which coincides with the appearance of fibronectin at the apical surface (see Figure 2C). To determine if these conditioned buffers were able to activate the TGF-β pathway in Calu-3 cells, we exposed CTL cells to them and phosphorylation of Smad2/Smad3 was monitored by western blot. As expected, treatment with buffers from polarized CFTR KD cells increased phosphorylation of Smad2/Smad3 as compared to cells treated with CTL buffers (p<0.01; Figures 3D, E). Moreover, we observed an overexpression of *Smad2* mRNA at D0, D3, and D14 of ALI (p<0.05; Figure S2B) and protein at D14 of ALI culture (p<0.001; Figure S2C, D). Smad7 acts a negative regulator of the TGF-β pathway and as a target gene of TGF-β1 signaling ^36^. We found an opposite relationship between CTL and CFTR KD cells in terms of Smad7 protein expression. Smad7 is increased during ALI culture in CTL cells while it is decreased from D3 in CFTR KD cells, reaching significance at D14 (p<0.05; Figures S2E, F), suggesting a loss of the negative regulation. Thus, increased secretion and activation of TGF-β, as well as a reduction of negative regulation all indicate an increased activation of TGF-β signaling in the absence of CFTR in Calu-3 cells.

**Figure 3.**
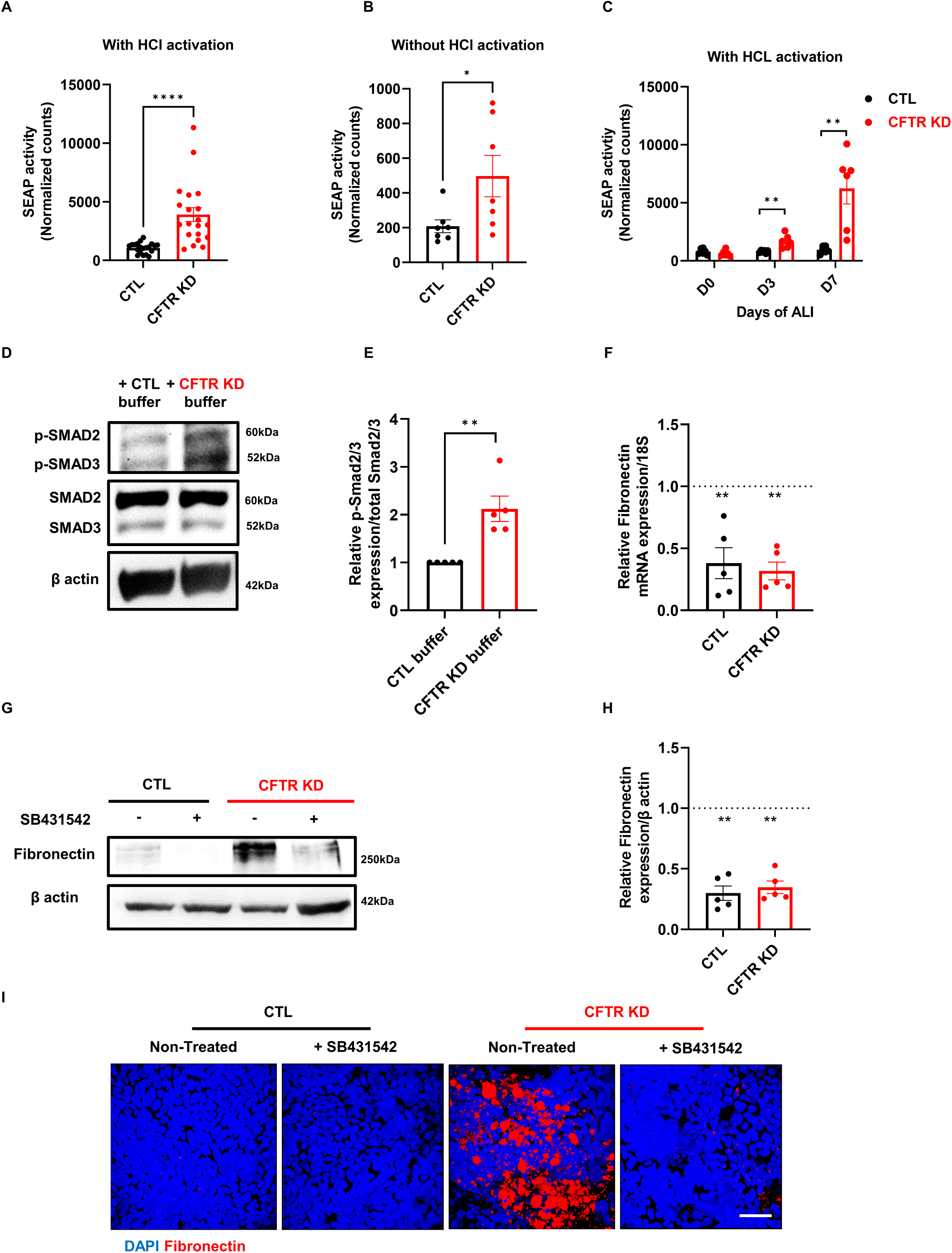
*CFTR* knockdown leads to over-production of TGF-β1 and over-stimulation of the TGF-β pathway. **(A and B)** SEAP activity of TGF-β reporter cell line after treatment with conditioned buffers for 24 hours from polarized CTL and CFTR KD cells with **(A)** and without **(B)** HCl activation (n=4). Mann-Whitney U test, * p<0.05, **** P<0.0001. (C) SEAP activity of TGF-β reporter cell line 24 hours after treatment with CTL and CFTR KD conditioned buffers during ALI with HCl activation (n=3). Mann-Whitney U test, ** P<0.01. **(D and E)** Representative western blot **(D)** and corresponding quantification **(E)** depicting phosphorylation of Smad2 (Ser465/467)/Smad3 (Ser423/425) and their total forms after treating CTL cells with CTL and CFTR KD conditioned buffers for 30 minutes (n=5). β-actin was used as a loading control. Mann-Whitney U test, ** p<0.01. **(F)** qPCR depicting mRNA expression of fibronectin in polarized CTL and CFTR KD cells after treatment with 10 µM SB431542 (n=5). 18S was used as an internal control. Mann-Whitney U test, ** p<0.01. **(G and H)** Representative western blot **(G)** and the corresponding quantification **(H)** depicting fibronectin protein expression in polarized CTL and CFTR KD cells after treatment with 10 µM SB431542 (n=5). β-actin was used as a loading control. Mann-Whitney U test, ** p<0.01. Dotted lines in **F** and **H** depict non-treated conditions. **(I)** Top view of 3D reconstructions of z-stack confocal images depicting fibronectin (red) and DAPI (blue) in polarized CTL and CFTR KD cells after treatment with 10 µM SB431542 (n=3). Scale bar: 40 µm. See also Figure S2.

Since we observed an apparent increase in TGF-β signaling, we sought to determine if pharmacological inhibition of the TGF-β pathway could prevent the overexpression and apical deposition of fibronectin in CFTR KD cells. To this end, we used a very potent and selective inhibitor of TGF-βRI, SB431542, from D7-D14 of ALI culture, thereby targeting the period during polarization that coincided with an increase in fibronectin expression in CFTR KD cells. SB431542 treatment led to a significant decrease in fibronectin mRNA expression both in CTL and CFTR KD cells (p<0.01; Figure 3F). This decrease in mRNA was paralleled with a decrease in fibronectin protein expression (p<0.01; Figures 3G, H). Next, we sought to determine if the inhibition of the TGF-β pathway could restore polarity by monitoring the deposition of apical fibronectin by immunostaining. Interestingly, SB431542 treatment fully prevented apical deposition of fibronectin in CFTR KD cells (Figure 3I). Altogether, these results indicate that the increased activation of the TGF-β pathway leads to the overexpression of fibronectin and its apical deposition in CFTR KD cells.

### *CFTR* knockdown leads to DKK1 over-secretion and inactivation of canonical Wnt signaling in Calu-3 cells

It is known that TGF-β signaling crosstalks with the Wnt pathway ^37^. Interestingly, DKK1, an inhibitor of the Wnt signaling pathway, was predicted by GSEA of the RNA-seq to be significantly enriched (Figure 1A), which was confirmed by RNAscope® (Figure 1C) and at the protein level in CF epithelia (Figure 1F, G). We then investigated if this was reproducible in the Calu-3 cell model. Indeed, KD of *CFTR* was associated with increased *DKK1* mRNA expression during ALI culture (p<0.05; Figure 4A). Next, we incubated polarized Calu-3 cells with a physiological saline buffer for different time points to collect apical secretions. Western blot was then performed to monitor DKK1 secretion. Already after one hour post-incubation, an increased secretion of DKK1 was observed in CFTR KD conditioned buffer, and this continued until 48 hours post-incubation (Figure 4B). Furthermore, this increased secretion could be observed as early as D3 of ALI culture and accumulated until D7 (Figure 4C).

**Figure 4.**
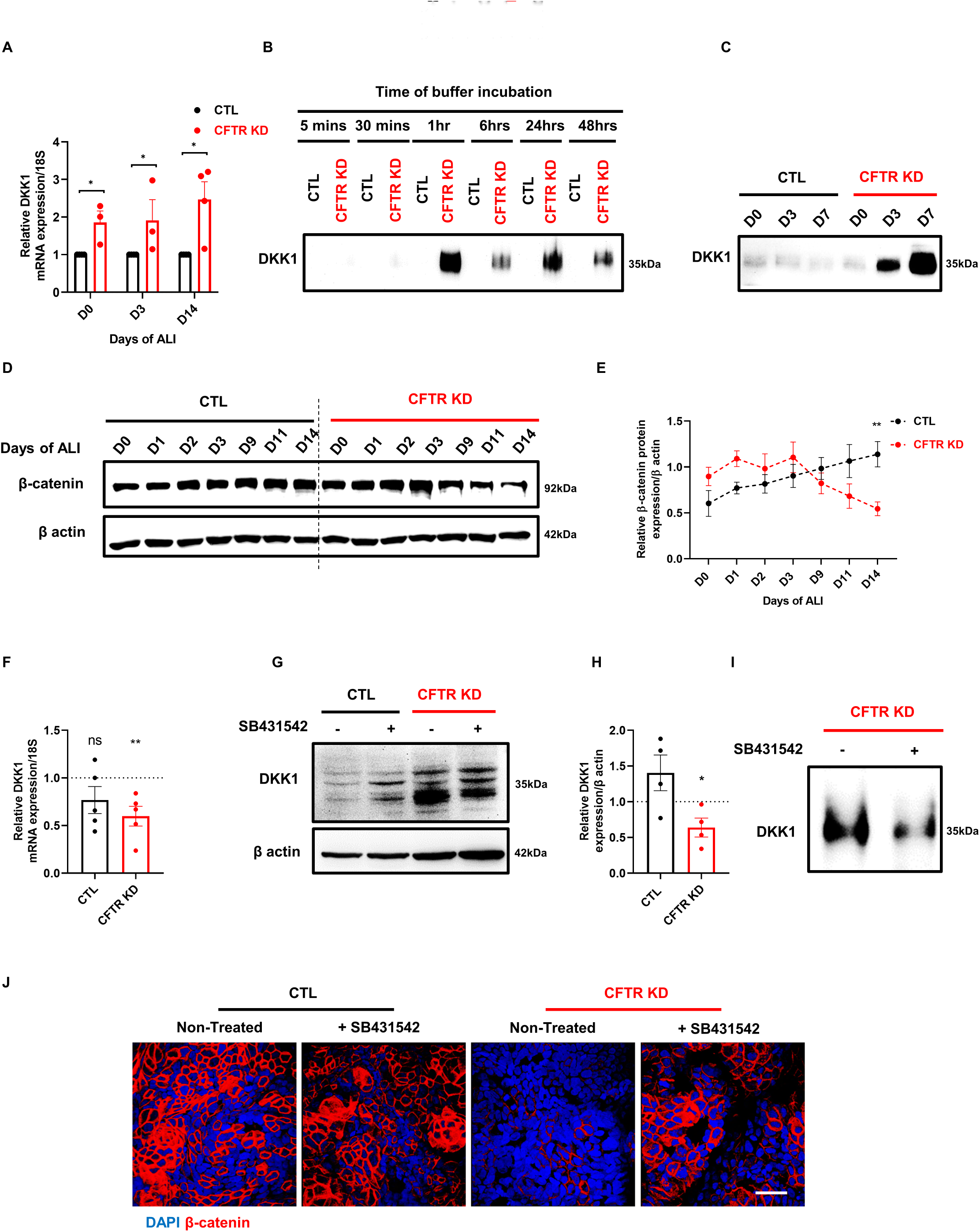
*CFTR* knockdown leads to DKK1 over-secretion and inactivation of canonical Wnt signaling in Calu-3 cells. **(A)** qPCR depicting DKK1 mRNA expression in CTL and CFTR KD cells during ALI (n=3). 18S was used as an internal control. Mann-Whitney U test, * p<0.05. **(B)** Representative western blot depicting DKK1 secretions in polarized CTL and CFTR KD cells, after incubation with buffers for different time points (n=3). **(C)** Representative western blot depicting DKK1 secretions in CTL and CFTR KD cells during ALI after incubation with buffer for 24 hours (n=3). **(D and E)** Representative western blot **(D)** and the corresponding quantification **(E)** depicting β-catenin protein expression in CTL and CFTR KD during ALI (n=6). β-actin was used as a loading control. Mann-Whitney U test, ** p<0.01. **(F)** qPCR depicting DKK1 mRNA expression in polarized CTL and CFTR KD cells after treatment with 10 µM SB431542 (n=5). 18S was used as an internal control. Mann-Whitney U test, ** p<0.01. **(G and F)** Representative western blot **(G)** and the corresponding quantification **(F)** depicting DKK1 protein expression in polarized CTL and CFTR KD cells after treatment with 10 µM SB431542 (n=4). β-actin was used as a loading control. Mann-Whitney U test, * p<0.05. Dotted lines in **F** and **H** depict non-treated conditions. **(I)** Representative western blot depicting DKK1 secretions in polarized CFTR KD cells after treatment with 10 µM SB431542, after 24 hours of buffer incubation (n=3). **(J)** Top view of 3D reconstructions of z-stack confocal images depicting β-catenin (red) and DAPI (blue) in polarized CTL and CFTR KD cells after treatment with 10 µM SB431542 (n=2). Scale bar: 40 µm. See also Figure S3.

Secreted DKK1 inhibits the canonical Wnt pathway by degrading β-catenin, the main signal transducer of the pathway ^17^. Therefore, we investigated whether the overexpression and over-secretion of DKK1 correlated with an inactivation of the Wnt pathway by monitoring the expression of β-catenin. β-catenin protein expression was similar between CTL and CFTR KD cells from D0 to D3 of ALI culture (Figures 4D, E). However, from day 9 (D9), β-catenin expression began to decrease until significantly at D14 (p<0.01; Figures 4D, E), correlating with the accumulation of DKK1 secretion in CFTR KD cells. In addition, we performed immunostaining of total β-catenin during ALI and z-stack images indicate that membrane localized β-catenin decreases from D9 of ALI until D14 in CFTR KD cells (Figure S3A). Moreover, we performed immunodetection of nuclear β-catenin with an antibody recognizing phosphorylated serine 552. Immunostaining experiments revealed a decrease in nuclear β-catenin in CFTR KD cells at D3, whereas the CTL cells continued to express this form until D14, suggesting decreased Wnt signaling in CFTR KD cells (Figure S3B). Next, we monitored β-catenin degradation, as a cause for decreased expression, in both CTL and CFTR KD cells at D14 of ALI culture using cycloheximide (CHX) chase assay. Indeed, after 24 hours of CHX treatment, β-catenin expression was strongly decreased in CFTR KD cells as compared to 0 hours treatment (p<0.01; Figures S3C, D). CTL cells on the other hand had a stable expression of β-catenin (Figures S3C, D). These results suggest that β-catenin phosphorylation and stability are decreased in CFTR KD cells, which is potentially caused by the increased secretion of DKK1 and inhibition of the canonical Wnt pathway.

Next, we investigated a potential link between Wnt and TGF-β pathways by monitoring DKK1 expression and secretion in CTL and CFTR KD cells after pharmacological inhibition of the TGF-β pathway. Treatment with SB431542 led to a small but significant decrease in DKK1 mRNA and protein expression (p<0.01, p<0.05; Figures 4F-H), as well as a lower secretion in CFTR KD cells (Figure 4I). This decrease in DKK1 secretion was associated with an increase in membrane-bound β-catenin in CFTR KD cells, as shown by confocal microscopy (Figure 4J). These results suggest an aberrant involvement of the TGF-β pathway in the modulation of the Wnt pathway during the establishment of epithelial polarization in CFTR KD cells.

### Pharmacological activation of Wnt signaling prevents fibronectin deposition

We next sought to determine if the loss of β-catenin protein expression and thus inactivation of the Wnt pathway was associated with the overexpression of fibronectin in CFTR KD cells. To this end, we treated Calu-3 cells with CHIR99021, a small molecule Wnt activator that blocks the activity of GSK-3β, thereby preventing the degradation of β-catenin. Treatment with CHIR99021 for 48 hours at D12-D14 of ALI culture, thus targeting β-catenin expression when it is significantly decreased in CFTR KD cells, was indeed able to increase total β-catenin expression in both CTL and CFTR KD cells (p<0.001; Figures 5A, B), without affecting its mRNA expression (Figure 5C). Furthermore, confocal images revealed an increase in membrane-bound β catenin in both CTL and CFTR KD cells (Figure 5D). Interestingly, total fibronectin was significantly reduced in CFTR KD cells (p<0.001; Figures 5E, F), whilst unchanged in CTL cells (Figures 5E, F). Fibronectin mRNA expression was also decreased after CHIR99021 treatment (p<0.05; Figure 5G). Moreover, we monitored apical deposition of fibronectin in CFTR KD cells by confocal microscopy and this was completely prevented after CHIR99021 treatment, thereby restoring polarity in CFTR KD cells (Figure 5H). Thus, activation of the Wnt pathway counteracts the TGF-β1-induced fibronectin overexpression in CFTR KD cells. Overall, these results suggest a complex interplay between the TGF-β and Wnt pathways in regulating fibronectin expression, which is dysregulated in CF HAECs.

**Figure 5.**
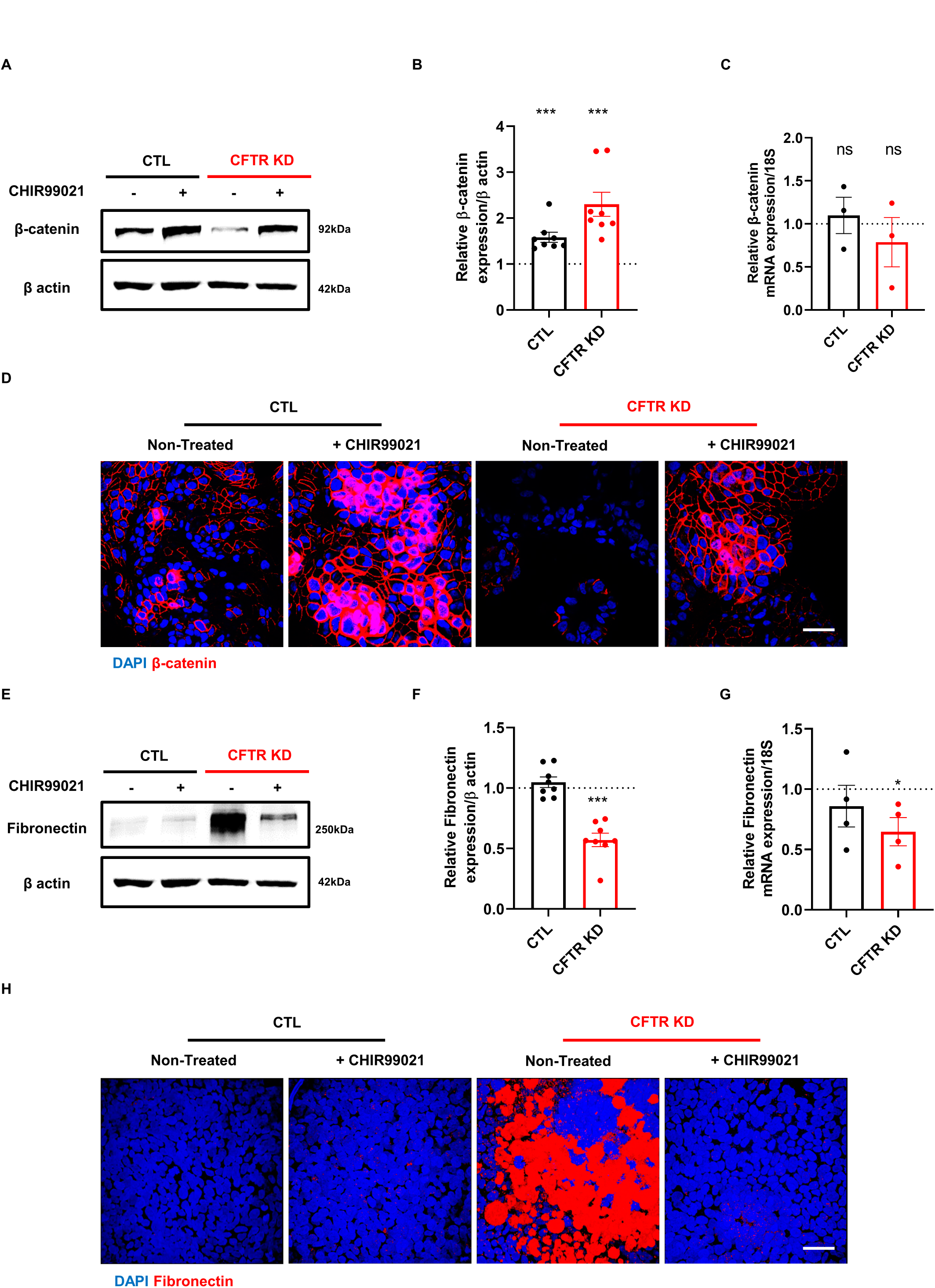
Pharmacological activation of Wnt signaling prevents fibronectin deposition. **(A and B)** Representative western blot **(A)** and corresponding quantification **(B)** depicting β-catenin protein expression in polarized CTL and CFTR KD cells after treatment with 10 µM CHIR99021 (n=8). β-actin was used as a loading control. Mann-Whitney U test, *** p<0.001. **(C)** qPCR depicting β-catenin mRNA expression in polarized CTL and CFTR KD cells after treatment with 10 µM CHIR99021 (n=3). 18S was used as an internal control. **(D)** Confocal images depicting β-catenin (red) and DAPI (blue) at the apical side of polarized CTL and CFTR KD cells (n=1). **(E and F)** Representative western blot **(E)** and corresponding quantification **(F)** depicting fibronectin protein expression in polarized CTL and CFTR KD cells after treatment with 10 µM CHIR99021 (n=8). β-actin was used as a loading control (same blot as in **A**). Mann-Whitney U test, *** p<0.001. **(G)** qPCR depicting fibronectin mRNA expression in polarized CTL and CFTR KD cells after treatment with 10 µM CHIR99021 (n=4). 18S was used as an internal control. Mann-Whitney U test, * p<0.05. Dotted lines in **B, C, F** and **G** depict non-treated conditions. **(H)** Top view of 3D reconstructions of z-stack confocal images depicting β-catenin (red) and DAPI (blue) in polarized CTL and CFTR KD cells after treatment with 10 µM CHIR99021 (n=3) Scale bars: 50 µm (upper panel), 40 µm (lower panel).

### Akt pathway modulates TGF-β signaling and DKK1 secretion in CFTR KD cells

A regulatory loop between Akt and GSK-3β is well established ^38, 39^. Indeed, CHIR99021 prevented the phosphorylation of Akt in both CTL and CFTR KD cells (p<0.001, p<0.0001; Figure S4A, B), confirming that GSK-3β activates Akt signaling under normal circumstances. To determine if Akt signaling interferes with TGF-β and Wnt pathways, we treated polarized Calu-3 cells with an allosteric, highly selective inhibitor of pan-Akt, MK2206. First, we monitored the expression of total and active, phosphorylated Akt (p-Akt) by western blot. Interestingly, we found that the p-Akt/Akt ratio was increased due to decreased expression of total Akt in CFTR KD cells (p<0.01; Figure S4C, D), validating the need to inhibit the pathway. We then confirmed that MK2206 treatment dampened Akt signaling without affecting total Akt expression in both CTL and CFTR KD cells (p<0.05; Figure S4E-G).

To better understand the interplay between Akt, TGF-β and Wnt pathways in the regulation of fibronectin expression, we first monitored the phosphorylation of Smad2 after MK2206 treatment, and surprisingly, inhibition of the Akt pathway decreased the phosphorylation of Smad2 in CFTR KD cells only (Figure S4H). Furthermore, TGF-β reporter cell line was treated with buffers collected from polarized Calu-3 cells after treatment with MK2206 and revealed reduced secretion of total TGF-β in the CFTR KD cells (p<0.01; Figure 6A), and in the CTL cells albeit to a lesser extent (p<0.05; Figure 6A). Because the TGF-β pathway was found to modulate DKK1 expression in CFTR KD cells, we also verified the effect of Akt inhibition on DKK1 secretion. Consistent with its effect on Smad2 phosphorylation, MK2206 treatment reduced DKK1 secretion in CFTR KD cells (Figure 6B). Finally, we monitored fibronectin expression and apical deposition after inhibition of the Akt pathway. MK2206 treatment reduced fibronectin mRNA (p<0.01; Figure 6C) and protein expression in CFTR KD cells but not in CTL cells (p<0.05; Figures 6D, E). Furthermore, we observed a drastic loss of apical fibronectin deposition in CFTR KD cells after treatment (Figure 6F), suggesting that Akt signaling inhibition restored polarity of CFTR KD cells.

**Figure 6.**
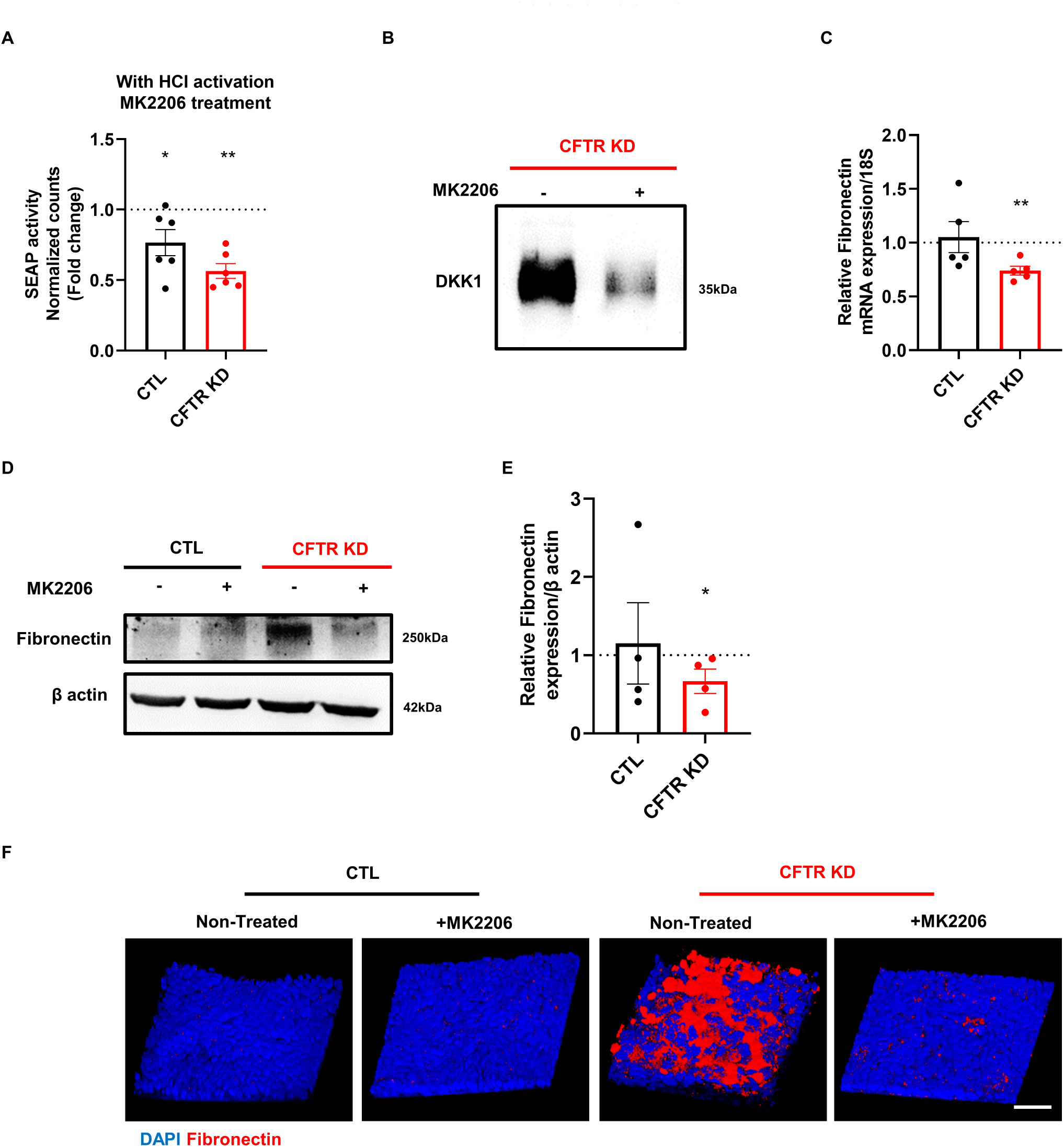
Akt pathway modulates TGF-β signaling and DKK1 secretion in CFTR KD cells. **(A)** SEAP activity of TGF-β reporter cell line after treatment with conditioned buffers from polarized CTL and CFTR KD cells with HCl activation, after treatment with 5 µM MK2206 (n=3). Mann-Whitney U test, * p<0.05, ** p<0.01. **(B)** Representative western blot depicting DKK1 secretions in polarized CFTR KD cells after treatment with 5 µM MK2206, after 24 hours of buffer incubation (n=3). **(C)** qPCR depicting fibronectin mRNA expression in polarized CTL and CFTR KD cells after treatment with 5 µM MK2206 (n=5). 18S was used as an internal control. Mann-Whitney U test, ** p<0.01. **(D and E)** Representative western blot **(D)** and corresponding quantification **(E)** depicting fibronectin protein expression in polarized CTL and CFTR KD cells after treatment with 5 µM MK2206 (n=4). β-actin was used as a loading control (same as in **A**). Mann-Whitney U test, * p<0.05. Dotted lines in **A, C and E** depict non-treated conditions. **(F)** 3D reconstructions of z-stack confocal images depicting fibronectin (red) and DAPI (blue) in polarized CTL and CFTR KD cells after treatment with 5 µM MK2206 (n=3). Scale bar: 40 µm. See also Figure S4.

These results indicate an imbalance between the TGF-β, Wnt and Akt signaling pathways with over-secretion of TGF-β1 and DKK1 in CFTR KD cells. This leads to an over-production and apical deposition of fibronectin and thus an altered polarity of the CF epithelium.

### Pharmacological modulation of TGF-β, Wnt or Akt signaling pathways prevented fibronectin apical deposition in primary CF HAECs

To validate the results obtained in Calu-3 cells, we monitored fibronectin expression after treatment with the pharmacological drugs in well-differentiated primary HAECs. First, confocal microscopy confirmed an overexpression and apical deposition of fibronectin in CF HAECs (p<0.0001; Figure 7A, B). Interestingly, fibronectin expression and apical deposition were decreased after activating the Wnt pathway, after blocking the Akt pathway and after blocking the TGF-β pathway, with MK2206 and SB431542 treatments exhibiting the strongest effects (p<0.05, p<0.0001, p<0.0001; Figures 7A, B). Next, β-catenin expression was monitored and confocal microscopy revealed a decrease in CF HAECs and this was confirmed after quantification of the staining (p<0.01; Figures 7A, C). We then monitored β-catenin expression after treatment with the pharmacological drugs. Interestingly, all the treatments led to an increase in membrane-bound β-catenin expression with CHIR99021 and SB431542 treatments displaying the strongest effects (p<0.0001, p<0.01, p<0.0001; Figures 7A, C). These results indicate that the unbalanced TGF-β, Wnt and Akt signaling pathways in CF HAECs lead to loss of apicobasal polarity associated with β-catenin degradation and ectopic accumulation of fibronectin on the epithelium surface.

**Figure 7.**
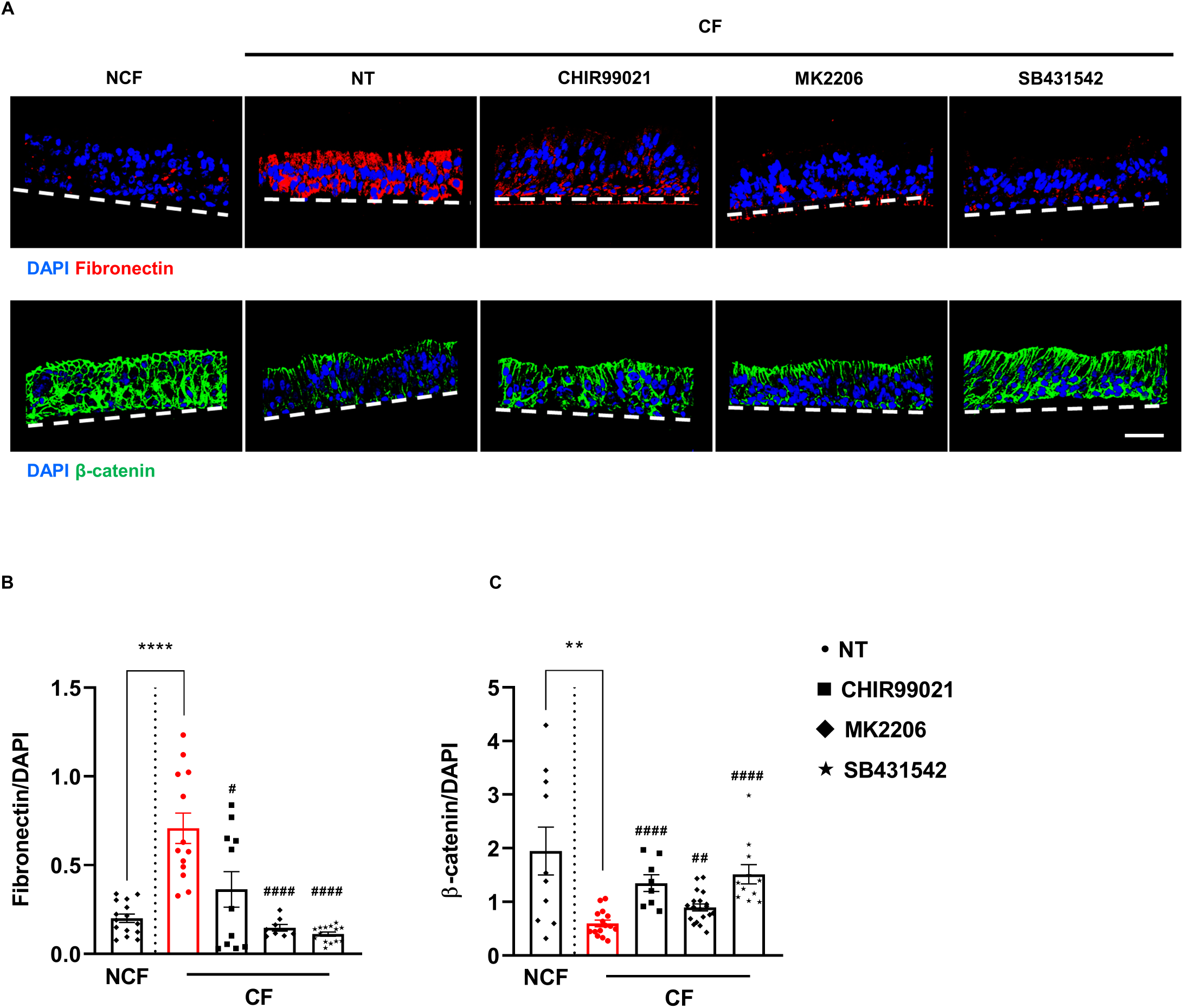
Pharmacological modulation of TGF-β, Wnt or Akt signaling pathways prevented fibronectin apical deposition in primary CF HAECs. **(A, B and C)** 3D reconstructions of z-stack confocal images depicting fibronectin (red), and β-catenin (green) and DAPI (blue) **(A)** and the corresponding quantifications (**B**, fibronectin, **C**, β-catenin) after treatment with 10 µM CHIR99021, 5 µM MK2206 and 10 µM SB431542, in differentiated CF (N=6) HAECs. NCF is shown as a comparison (N=4). Unpaired 2-tailed Student’s t test, ** p<0.01, **** p<0.0001, ^#^ p<0.05, ^##^ p<0.01, ^####^ p<0.0001. * Depicts comparison between NCF and CF, ^#^ depicts comparison between CF non-treated and treated conditions. Dashed lines in A depict basal side. Scale bar: 40 µm.

## DISCUSSION

Airway epithelium barrier integrity relies on the development of an apicobasal polarity of epithelial cells. As the airway epithelium is frequently exposed to injury, its regeneration is critical to restore integrity and barrier function. Epithelial repair is a multi-step process, the regulation of which remains poorly understood, involving a dynamic balance between migration, proliferation and restoration of cell polarity. In CF, increasing evidence points to an apicobasal polarity defect of the airway epithelium which may contribute to the chronic infection that characterizes this respiratory disease. In this study, we report an uncoordinated interplay between TGF-β, Wnt and Akt signaling that causes β-catenin degradation and surface accumulation of fibronectin during epithelial regeneration.

Combining RNA-seq GSEA, RNA *in situ* hybridization, reporter cell line assays and immunoblot, we found that TGF-β signaling was altered during the repair process of primary CF HAECs. Fully differentiated primary CF HAECs displayed increased TGF-β expression and secretion. This observation was confirmed in polarizing Calu-3 cell line after knockdown of *CFTR*, indicating the causal relationship between the CFTR defect and abnormal regulation of this protein. Importantly, we observed that TGF-β1 secretion is increased at the apical surface of the CF epithelia. Airway epithelial polarity can impact on TGF-β1 signaling and the subsequent production of fibronectin, as previously reported ^40, 41^. Among the mechanisms known to regulate TGF-β1 secretion and activation, which include, low pH, reactive oxygen species (ROS), inflammation, proteolysis and integrins ^15, 42, 43^, elevated inflammation and ROS production are characteristics of the CF phenotype ^44, 45^, Also, it has been reported that the expression of furin, the convertase mediating cleavage of latent TGF-β1 which is necessary for its subsequent activation ^46^, is elevated in the IB3-1 CF bronchial cell line 17948127 ^47^. In addition to enhanced secretion of bioactive TGF-β1, we also reported increased expression and phosphorylation of Smad2 but decreased expression of Smad7, which may thus provide a positive loop for activation of the TGF-β pathway and its target genes, including fibronectin ^35^. In CFTR KD cells, induction of fibronectin expression was indeed prevented at transcriptional and protein levels by the TGF-βR1 inhibitor SB431542. Aberrant TGF-β activity in CF has been suggested in other studies ^48, 49^. Importantly a study showed that TGF-β1 impaired CFTR biogenesis and function after correction with CFTR modulators in CF bronchial epithelial cells ^50^. Finally, genetic variations in the 5’ end of the TGF-β1 gene have been identified and shown to contribute to CF disease severity ^25, 51^. Thus, these studies further corroborate our findings and highlight the importance of TGF-β research as a complementary strategy to CFTR modulators in CF ^52^.

Not only was TGF-β signalling increased, we also report for the first time inactivation of the Wnt pathway via increased apical secretion of DKK1 in primary CF and CFTR KD HAECs. DKK1 protein is an endogenous inhibitor of the canonical Wnt signaling pathway, stimulating the degradation of β-catenin through GSK-3β ^17^. In CFTR KD cells, immunoblotting and immunofluorescence of β-catenin and CHX chase assay revealed that an over-secretion of DKK1 is indeed associated with inhibition of the Wnt pathway. In addition, DKK1 mRNA and protein expression were reduced by SB431542, indicating that TGF-β1-induced DKK1 expression and secretion may be responsible for the inhibition of the Wnt pathway in CFTR KD cells. Importantly, reactivating Wnt signaling in CFTR KD cells with the GSK-3β inhibitor CHIR99021 restored β-catenin pools and its translocation to the nucleus, and prevented the mRNA and protein overexpression of fibronectin. This observation further indicates that β-catenin stability is decreased in CFTR KD cells. β-catenin is both a component of adherens junctions, linking cadherin to the actin cytoskeleton, and a signaling molecule, thus its destabilization may explain the impairment in proliferation and delayed differentiation during the repair process of CF epithelia ^53–56^. It is worth noting a study showing the regulation of Wnt/β-catenin by CFTR during lung development ^57^. In addition, CFTR-β-catenin interaction has been reported in non-airway CF tissues ^58–60^. A peak in cell proliferation followed by a gradual decrease usually precedes polarization and differentiation of airway epithelial cells ^61, 62^. In this context, Wnt and TGF-β pathways can regulate each other in a complex manner, with both positive and negative feedback loops existing between the two pathways ^63^. Thus, the disrupted interplay between TGF-β and Wnt signaling pathways may explain in part the strong inhibition of cell proliferation and the abnormal deposition of fibronectin at the luminal side of the epithelium, leading to apicobasal polarity defect in CFTR KD cells.

Fibronectin and its receptor β1-integrin have been shown to be transiently overexpressed and apically localized in repairing airways leading to increased *Pa* adhesion to the epithelium ^64, 65^. This process has also been associated with dedifferentiation of the epithelium and loss of polarity ^64, 65^. Loss of epithelial polarity and impairment in proliferation are also key hallmarks of EMT ^12, 13^. Notably, the TGF-β and Wnt pathways have been implicated in the EMT process in CF ^58, 66–68^. Altogether, our findings, along with these studies, indicate that temporal dysregulation of TGF-β and Wnt pathways leads to a partial EMT phenotype of the CF airway epithelium.

Akt is another key interactor of the Wnt pathway through the phosphorylation of its regulatory sites by GSK-3β ^38^. Aligned with this idea, CHIR99021 fully inhibited phosphorylation of Akt in both CTL and CFTR KD cells. Thus, DKK1-mediated inhibition of the Wnt pathway in CFTR KD cells could also affect Akt activation and downstream protein stability. Surprisingly, inhibition of Akt activity with MK2206 decreased p-Smad2/3 only in CFTR KD cells, reducing transcriptional and protein expression of fibronectin as well as its accumulation on the apical surface. These suggest that an aberrant crosstalk between Akt and TGF-β pathways is specifically occurring during the polarization process of CFTR-defective epithelial cells. Interestingly, it has been shown that α5β1 integrin clusters to the cell surface can immobilize fibronectin, leading to increased Akt phosphorylation and activation ^69^. Others and we have previously reported overexpression of β1-integrin at the surface of CF HAECs ^10, 11, 34^. Therefore, the presence of apical fibronectin could serve as a positive loop and lead to further activation of Akt in CF cells. Thus, modulating Akt pathway would be critical in the modulation of both TGF-β and Wnt signaling pathways ^30^. Consistent with this idea, MK2206 reduced TGF-β1 and DKK1 secretion in CFTR KD cells. It is widely believed that PI3K/Akt signaling enhances Wnt signals by modulating transcriptional activity and stability of β-catenin ^70^. Thus, aberrant PI3K/Akt activity can lead to alterations in the regulation of multiple cell processes, including survival, proliferation, polarization and metabolism of CF HAECs. Whether TGF-β and Wnt non-canonical pathways are also altered by the aberrant PI3K/Akt activity in CF HAECs remains to be investigated.

It is remarkable how primary HAECs from donors carrying different *CFTR* mutations were all associated with an abnormal interplay between TGF-β, Wnt and Akt pathways. Importantly, pharmacological activation of the Wnt pathway or inhibition of TGF-β and Akt pathways, restored β-catenin expression and prevented fibronectin deposition at the apical surface of primary CF HAECs. Combinatorial treatments with the different pharmacological drugs could potentially further augment their effects in restoring polarity. Indeed, drug combinations at low doses could potentially maximize efficacy, lead to lower toxicity and reduce compensatory effects thus enhancing therapeutic potential ^71^. Ultimately, this would abolish apical fibronectin deposition in CF models exhibiting different classes of *CFTR* mutations, restore apicobasal polarity and prevent bacterial infections ^10, 34^. Moreover, CFTR modulators have been shown to be unable to completely eradicate pathogens in CF airways ^72^. Thus, the underlying signals regulating polarization investigated in this study may shed light on potential complementary therapeutic targets for CF and other chronic airway diseases involving dysregulation of these pathways and defects in epithelial structural integrity.

## MATERIALS AND METHODS

### Cell culture and treatments

Primary HAECs were purchased from Epithelix Sàrl (Switzerland). They were isolated from bronchial biopsies and were cultured on Transwell inserts in MucilAir Culture Medium from Epithelix Sàrl and differentiated at the Air-liquid interface (ALI). The characteristics of the NCF and CF donors are provided in Supplemental Table 1. Calu-3 Human Airway Epithelial Cells (HAECs) (American Type Culture Collection, ATCC^®^ HTB-55^™^) were transfected with CRISPR-Cas9 lentiviral vector particles (GeneCopoeia™) targeting the *CFTR* gene to generate CFTR KD cells and its control (CTL), as previously described ^73^. They were cultured in Minimum Essential Medium (MEM) GlutaMax^™^ (ThermoFisher) supplemented with 10% heat-inactivated Fetal bovine Serum (FBS), 1% non-essential amino acids (NEAA) 100X, 1% HEPES 1M, 1% sodium pyruvate 100X, 0.25 μg/ml fungizone, 100 IU/ml penicillin and 100 μg/ml streptomycin in a humified incubator at 37°C with 5% CO2. Polarized Calu-3 cells were obtained by seeding 1.75 x 10^5^ cells on polyester membrane inserts (0.4μm pore size, 0.33 cm^2^ surface area, Costar^®^ Corning). At 100% confluency, the cells were grown at an Air-Liquid Interface (ALI) for 14 days.

Polarized Calu-3 cells were treated with 150 μg/ml cycloheximide (CHX, Sigma) in DMSO (Sigma, #41639), for 1, 2, 4, 6, and 24 hours at 37°C. Primary HAECs were treated with 10 µM CHIR99021 (Sigma), 10 µM SB431542 (Sigma), and 5 µM MK2206 (Selleckchem) in DMSO for 48 hours. Calu-3 cells were treated with the same conditions except with 10 µM SB431542 from D7 to D14 of ALI.

### RNA-seq

RNA was isolated and sequenced from differentiated Human Airway Epithelial Cells obtained from seven CF patients and six non-CF (NCF) patients as described previously_33._

### Gene set enrichment analyses

All annotated pathways for *Homo sapiens, Mus musculus, Rattus norvegicus, Danio rerio, Sus scrofa and Saccharomyces cerevisiae* available on WikiPathways database (http://www.wikipathways.org/index.php/WikiPathways) were used to create gene sets, and the KEGG metabolic pathways (KEGG http://www.genome.jp/kegg/) relative to GRCh38.80. Genes were ranked by their calculated fold-changes (Rank Metric Score, decreasing ranking). Gene set analysis was performed with the GSEA package Version 2.2 from the Broad Institute (MIT, Cambridge, MA) and was used to analyze the pattern of differential gene expression between the CF versus NCF groups. Gene set permutations were performed 1000 times for each analysis. The Normalized Enrichment Score (NES) was calculated for each gene set. GSEA results with a nominal FDR < 0.05 and abs(NES) > 1 were considered significant.

### RNAscope®

The RNAscope**®** assay was used to monitor TGF-β1 and DKK1 expression in primary HAECs. An RNAscope® Multiplex Fluorescent Reagent v2 Assay Kit (Advanced Cell Diagnostics, ACD) and highly sensitive probes for TGF-β1 and DKK1 (ACD) were used according to the manufacturer’s instructions. Briefly, FFPE sections were deparaffinized and pretreated with hydrogen peroxide (ACD) for 10 minutes, antigen retrieval was performed for 15 minutes at 100°C and protease pretreatment was performed for 15 minutes at 40°C. Next, the samples were incubated with probes targeting TGF-β1 and DKK1 for 2 hours at 40 °C in a HybEZ oven (ACD). Detection of hybridized complexes was performed with Opal^TM^ dyes (Akoya Biosciences) according to manufacturer’s instructions. Sections were counterstained with DAPI. Images were acquired with LSM700 confocal microscope and ZEN software (ZEISS). The images were analyzed using ZEN and ImageJ software.

### TGF-β bioassay

The TGF-β reporter cell line was a kind gift from Prof. Tony Wyss-Coray and was used as described in the protocol developed by Tesseur *et al.,* 2006 ^74^. Briefly, embryonic fibroblast from *TGF-β1^-/-^* mice were transfected with a reporter plasmid expressing TGF-β responsive Smad-binding elements coupled to secreted alkaline phosphatase (SEAP) reporter gene. TGF-β reporter cell line was cultured in Dulbecco′s Modified Eagle′s Medium (DMEM) high glucose (Sigma), supplemented with 10% heat inactivated FBS, 1% penicillin/streptomycin (P/S) and 100 μg/ml hygromycin B (Corning) in a humified incubator at 37°C with 5% CO_2_.

Apical surfaces of primary HAECs and Calu-3 cells were washed twice with PBS before incubation with 100 μl of physiological saline buffer (NaCl 154 mM, HEPES 10 mM, CaCl_2_ 1.2 mM) for 24 hours. TGF-β reporter cell line was seeded at 3 x 10^4^ cells per well in a 96-well plate overnight. The cells were then washed twice in PBS and incubated with 50 μl serum free DMEM medium supplemented with 1% P/S for 2 hours. 50 μl of conditioned buffers from primary HAECs and Calu-3 cells was added to TGF-β reporter cell line without activation or pre-activated by adding 0.2 μl of 1N HCl for 15 minutes, followed by neutralization by adding 0.2 μl of 1N NaOH at room temperature (RT). 10 ng/ml TGF-β1 (gift from Prof. B. Wehrle-Haller) was used as a positive control. 10 μl of supernatant was collected 24 hours after incubating TGF-β reporter cell line with conditioned buffers. SEAP activity was measured using Phospha-light^TM^ assay system (Thermofisher) according to manufacturer’s instructions with Hidex Sense Microplate Reader (Labgene).

### DKK1 secretion

Apical surfaces of primary HAECs and Calu-3 cells were washed twice with PBS before incubation with 20 μl of physiological saline buffer (NaCl 154 mM, HEPES 10 mM, CaCl_2_ 1.2 mM) for 24 hours or otherwise. 12.5 μl of Laemmli sample buffer (100 mM Tris-HCl (pH6.8), 100mM β-mercaptoethanol, 4% SDS, 0.2% Bromophenol blue, 20% glycerol) was added to conditioned buffers, heated for 10 minutes at 95°C and loaded onto 12% SDS polyacrylamide gels (Bio-Rad).

### RNA extraction, RT-PCR and qPCR

Total RNA was extracted from Calu-3 cells with RNeasy mini kit (Qiagen). Extracted RNA was quantified and purity was verified by Nanodrop 2000 spectrophotometer (ThermoFisher). gDNA wipeout buffer was used to remove Genomic DNA for 2 minutes at 42°C and cDNA was synthetized with the QuantiTect Reverse Transcription Kit (Qiagen). qPCR was performed with PowerUp SYBR Green Master Mix using the StepOnePlus Real-Time PCR system. All primers were purchased from Microsynth AG, and sequences are listed in Supplementary Table 2.

### Western blotting

Proteins were extracted from Calu-3 cells with a RIPA lysis buffer (150 mM sodium chloride, 50 mM Tris (pH 7.4), 1% NP-40 (Applichem), 0.1% SDS, 0.5% Sodium deoxycholate (DOC) and Roche cOmplete^TM^ Protease Inhibitor Cocktail (Roche)). Protein extracts were centrifuged at 14,000 x g at 4°C for 20 minutes and quantified with Pierce BCA Protein Assay Kit (ThermoFisher). 10 μg or 20 μg of proteins were separated by electrophoresis in denaturing SDS polyacrylamide gel (Bio-Rad). The proteins were then transferred onto Porablot NCP nitrocellulose membrane (Macherey-Nagel) and blocked for 1 hour at RT in 3% BSA (Applichem) in PBS-Tween buffer (0.01%) (ThermoFisher, Sigma). The membranes were then incubated overnight at 4°C with primary antibodies listed in the Supplemental table 3 with agitation. GAPDH and β-actin antibodies were used as loading controls. After primary antibody fixations, the membranes were rinsed with PBS-Tween buffer (0.01%) and incubated with horseradish peroxidase–coupled (HRP-coupled) secondary antibodies (Supplementary table 3). Finally, proteins were revealed by detection of HRP with the SuperSignal West Pico PLUS chemiluminescent substrate (ThermoFisher). The images were detected by the Syngene PXi image analysis system and analyzed using ImageJ software (NIH).

### Immunofluorescence

Primary HAECs were embedded in Tissue-Tek® O.C.T. compound and cryosections were immediately fixed in 4% paraformaldehyde solution (PFA) for 30 minutes at RT and permeabilized for 15 minutes at RT with 0.2% Triton 100 X buffer (Sigma). Calu-3 cells were fixed with 4% PFA for 20 minutes at RT and permeabilized for 15 minutes at RT with 0.2% Triton 100X buffer. Nonspecific sites were blocked with 1% BSA in PBS for 30 minutes at RT, and samples were then incubated with primary antibodies (Supplementary table 3) at 4°C overnight. After several PBS washes, target proteins were detected with secondary antibodies (Supplemental table 3) for 1 hour at RT. DAPI counterstaining was used to visualize the nuclei. The images and z-stacks were obtained with LSM700 confocal microscope and ZEN software (ZEISS). The images were analyzed using ZEN, and ImageJ (NIH).

### Statistical analysis

Values are represented as mean ± SEM. The symbol N represents the number of donors from which primary HAECs were isolated and the symbol n represents the number of replicates. The statistical tests were realized using GraphPad Prism 9.5. The differences between two groups were analyzed by Student’s *t*-test or the non-parametric Mann-Whitney test. The Two-Way analysis of variance (ANOVA) test was performed to examine the differences between more than two groups. p<0.05, p<0.01, p<0.001, p<0.0001 are considered significant and expressed as *, **, ***, ****, or ^#^, ^##^, ^###^, ^####^ respectively.

## AUTHOR CONTRIBUTIONS

M.B. and M.C.: conceived and planned the study design. T.I., M.Bacchetta. and M.B: carried out experiments. T.I., M.B., M.Bachmann, B.W-H and M.C.: participated to analyses and discussion. T.I., M.B. and M.C.: wrote the manuscript. M.C.: supervision, project administration and funding acquisition. All authors reviewed the manuscript.

## ACKNOWLEDGEMENTS

This work was supported by the Swiss National Science Foundation. We thank the Faculty of Medicine core facilities (University of Geneva): Bioimaging, Histology and Genomics Platform (iGE3). We also thank Marie-Thérèse A Bou Younes for performing initial Akt experiments, Sylvain Lemeille for Gene Set Enrichment Analysis, Prof. Doron Merkler and Ingrid Wagner for sharing RNAscope equipment and advice. We would also like to thank Juliette Simonin, Paul Bigot and Rana El Masri for their comments on the manuscript.

## DATA AVAILABILITY STATEMENT

The dataset for this article can be found at https://doi.org/

## SUPPLEMENTAL FIGURE LEGENDS

**Supplemental Figure 1. Schemes for key experiments on primary HAECs and Calu-3 cells. (A)** Reproducible circular wounds were applied to primary HAECs ALI cultures with an airbrush. NW = Non-wounded, W = wounded, pW = 24 hours post wound, WC = wound closure, pWC = 48 hours post wound closure. **(B)** Fully differentiated HAECs were incubated with a physiological saline buffer for 24 hours to collect secretions. **(C)** CTL and CFTR KD Calu-3 cells were cultured as monolayers on transwell filters until they were confluent before 14 days of ALI culture. Filters were obtained at different days of ALI. Proliferation/polarization balance is crucial during ALI.

**Supplemental Figure 2. C*F*TR knockdown leads to increased TGF-β signaling. (A and B)** qPCR depicting TGF-β1 **(A)** and Smad2 **(B)** mRNA expression in CTL and CFTR KD cells during ALI (n=3). 18S was used as an internal control. Mann-Whitney U test, * p<0.05. **(C and D)** Representative western blot **(C)** and the corresponding quantification **(D)** depicting Smad2 protein expression in CTL and CFTR KD during ALI (n=3). β-actin was used as a loading control. Mann-Whitney U test, *** p<0.001. **(E and F)** Representative western blot **(E)** and the corresponding quantification **(F)** depicting Smad7 protein expression in CTL and CFTR KD during ALI (n=6). β-actin was used as a loading control. Mann-Whitney U test, * p<0.05.

**Supplemental Figure 3. C*F*TR knockdown promotes β-catenin degradation an inhibition of Wnt signaling. (A)** 3D reconstructions of z-stack confocal images depicting β-catenin (red) and DAPI (blue) in CTL and CFTR KD cells during ALI (n=2). **(B)** Confocal images depicting phospho-β-catenin (S552) (red) and DAPI (blue) at the apical side of CTL and CFTR KD cells during ALI (n=2). **(C and D)** Representative western blot **(C)** and the corresponding quantification **(D)** depicting β-catenin protein expression in CTL and CFTR KD after treatment with cycloheximide for 24 hours (n=3). GAPDH was used as a loading control. Two-way Anova, ** p<0.01. Scale bars: 40 µm (upper panel), 50 µm (lower panel).

**Supplemental Figure 4. Akt signaling pathway interacts with the Wnt and TGF-β signaling pathways. (A and B)** Representative western blot **(A)** the corresponding quantification **(B)** depicting phosphorylation of Akt (Ser473) and its total form after treating polarized CTL and CFTR KD cells with 10 µM CHIR99021 (n=3). β-actin was used as a loading control. Unpaired 2-tailed Student’s t test, *** p<0.01, **** p<0.0001. **(C and D)** Western blot quantifications of total **(C)** and phospho-Akt (S473) **(D)** in polarized CTL and CFTR KD cells (n=6). β-actin was used as a loading control. Mann-Whitney U test, ** p<0.01. **(E, F and G)** Representative western blot **(E)** and corresponding quantifications depicting phosphorylation of Akt (Ser473) **(F)** and its total form **(G)** after treating polarized CTL and CFTR KD cells with 5 µM MK2206 (n=4). β-actin was used as a loading control. Mann-Whitney U test, * p<0.05. **(H)** Representative western blot depicting phosphorylation of Smad2 (Ser465/467) and its total form after treating polarized CTL and CFTR KD cells with 5 µM MK2206 (n=2). β-actin was used as a loading control. Dotted lines in **B, F** and **G** depict non-treated conditions.

**Supplementary Table 1. Clinicopathological characteristics of CF and NCF donors.**

^1^ Human Airway Epithelial Cells (HAECs).

^2^ TransEpithelial Electrical Resistance (TEER).

**Supplementary Table 2. Primer sequences used for qPCR experiments.**

**Supplementary Table 3. Key reagents and resources used in this study.**

