## Supplemental data for "Akt Drives TGF-β-induced Over-secretion of DKK1 and Impairment of Cystic Fibrosis Airway Epithelium Polarity"

Supplementary Figure 1

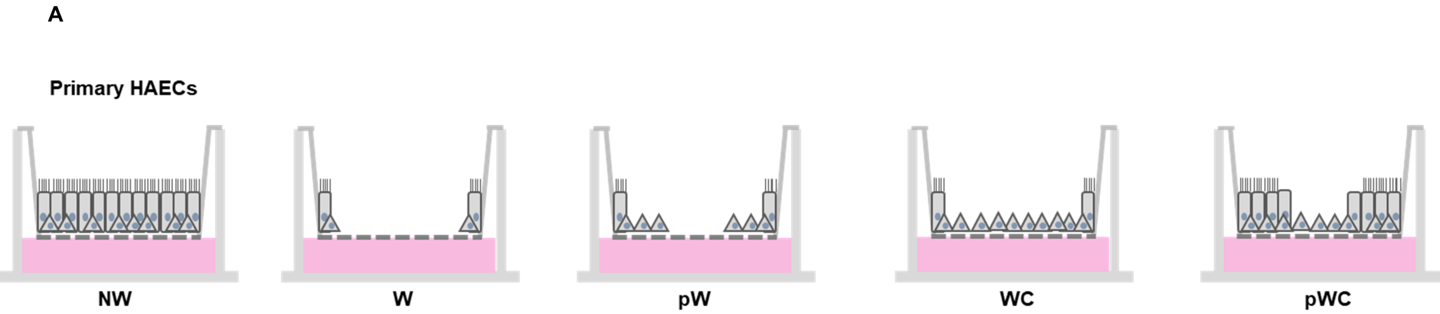


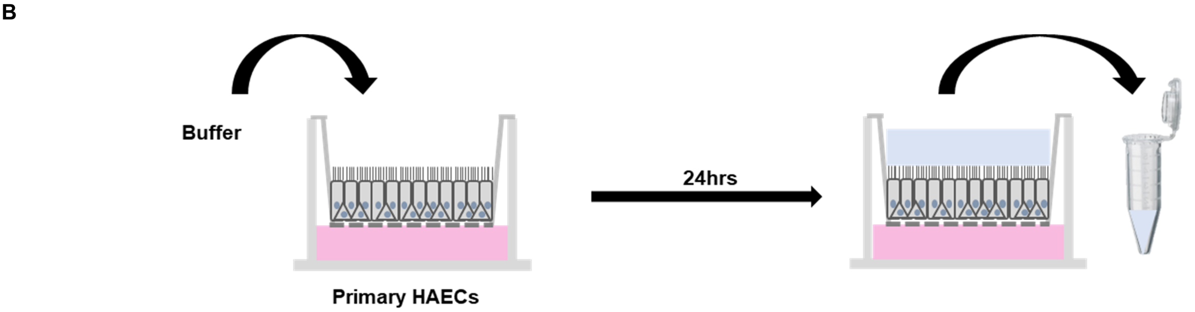


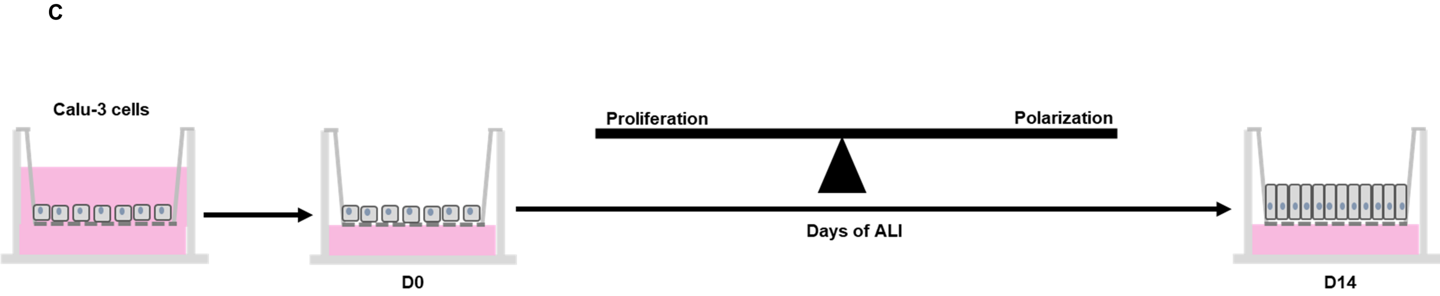

### Supplementary figure 2

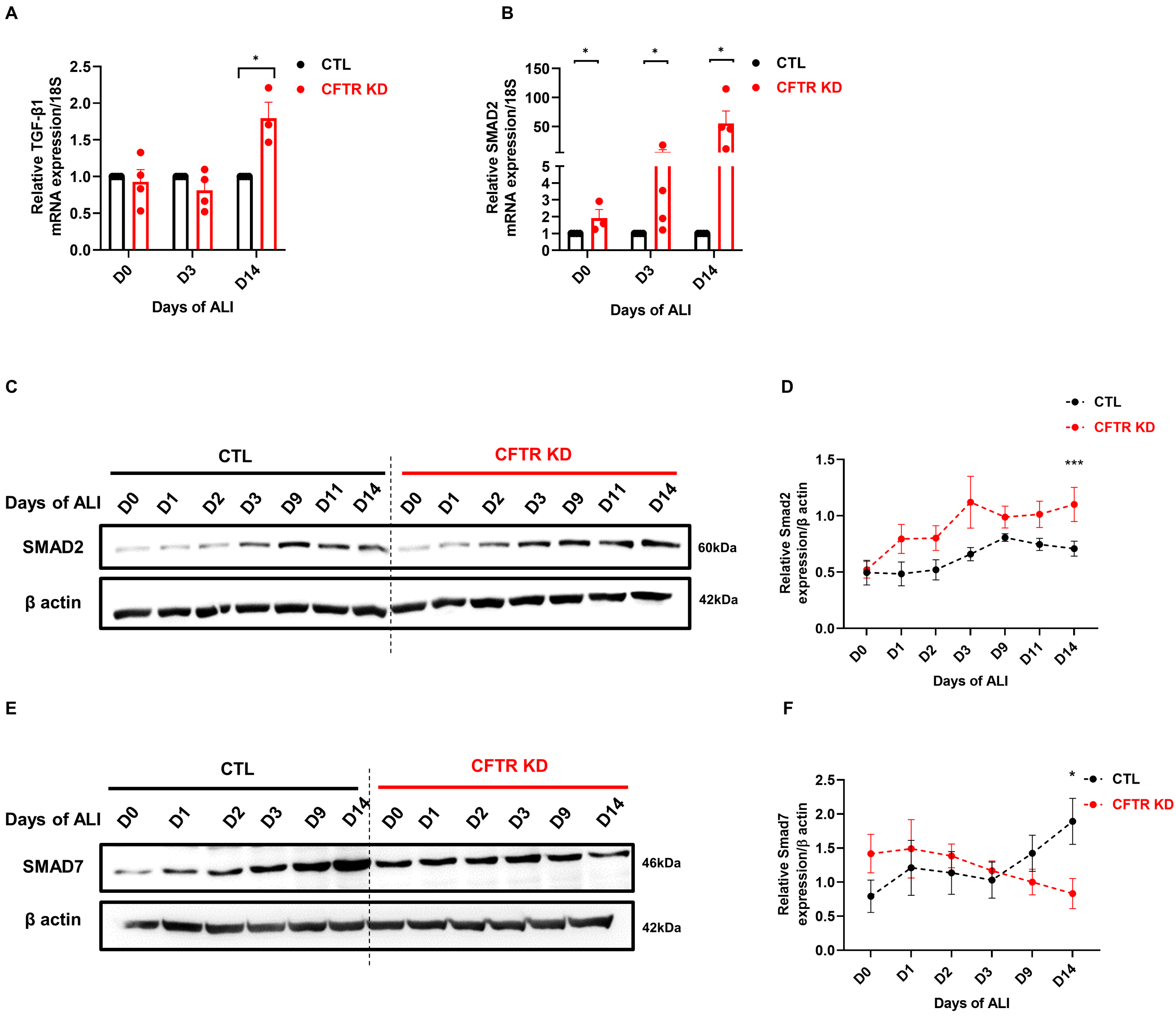

Supplementary figure 3

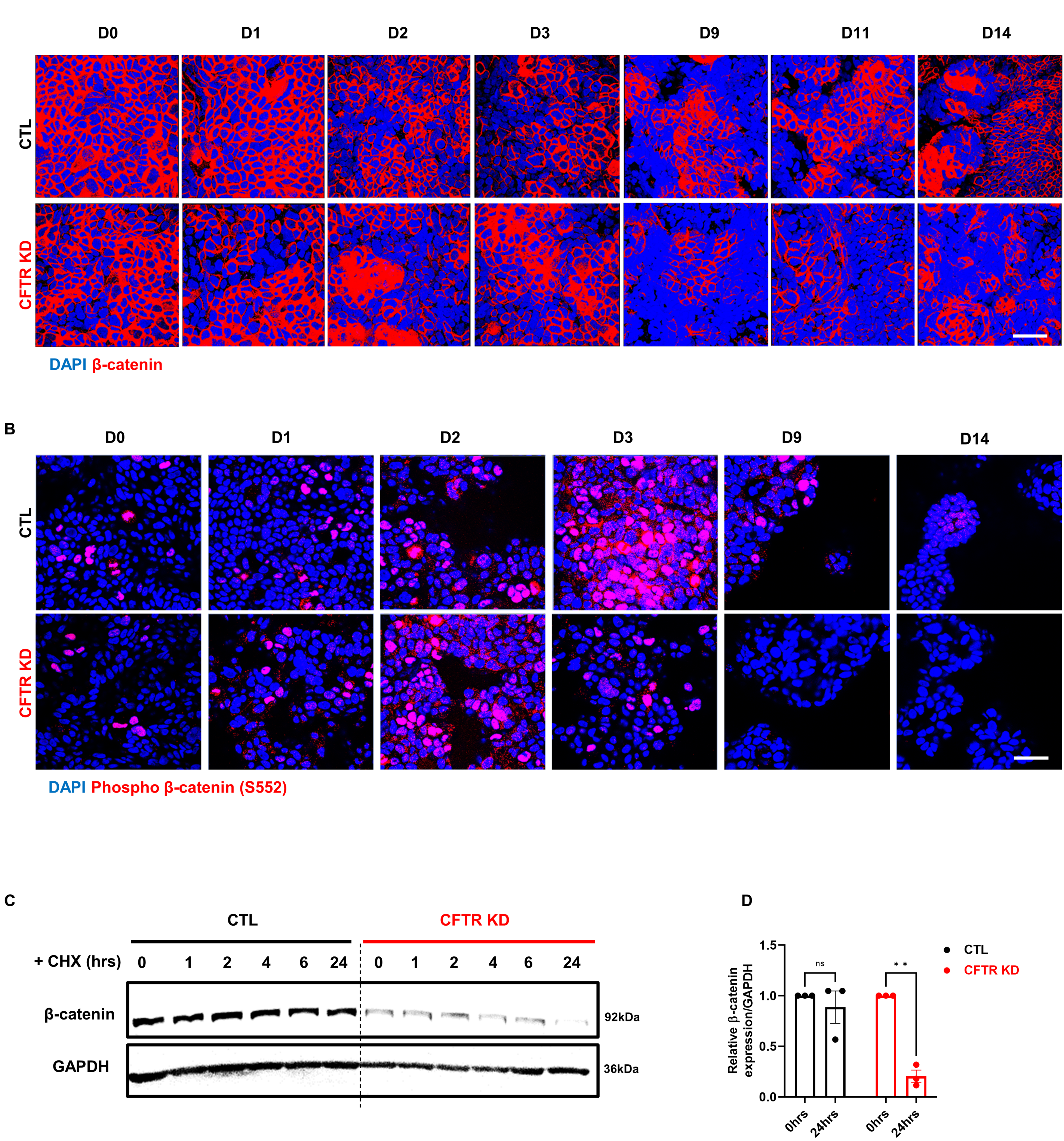

Supplementary figure 4

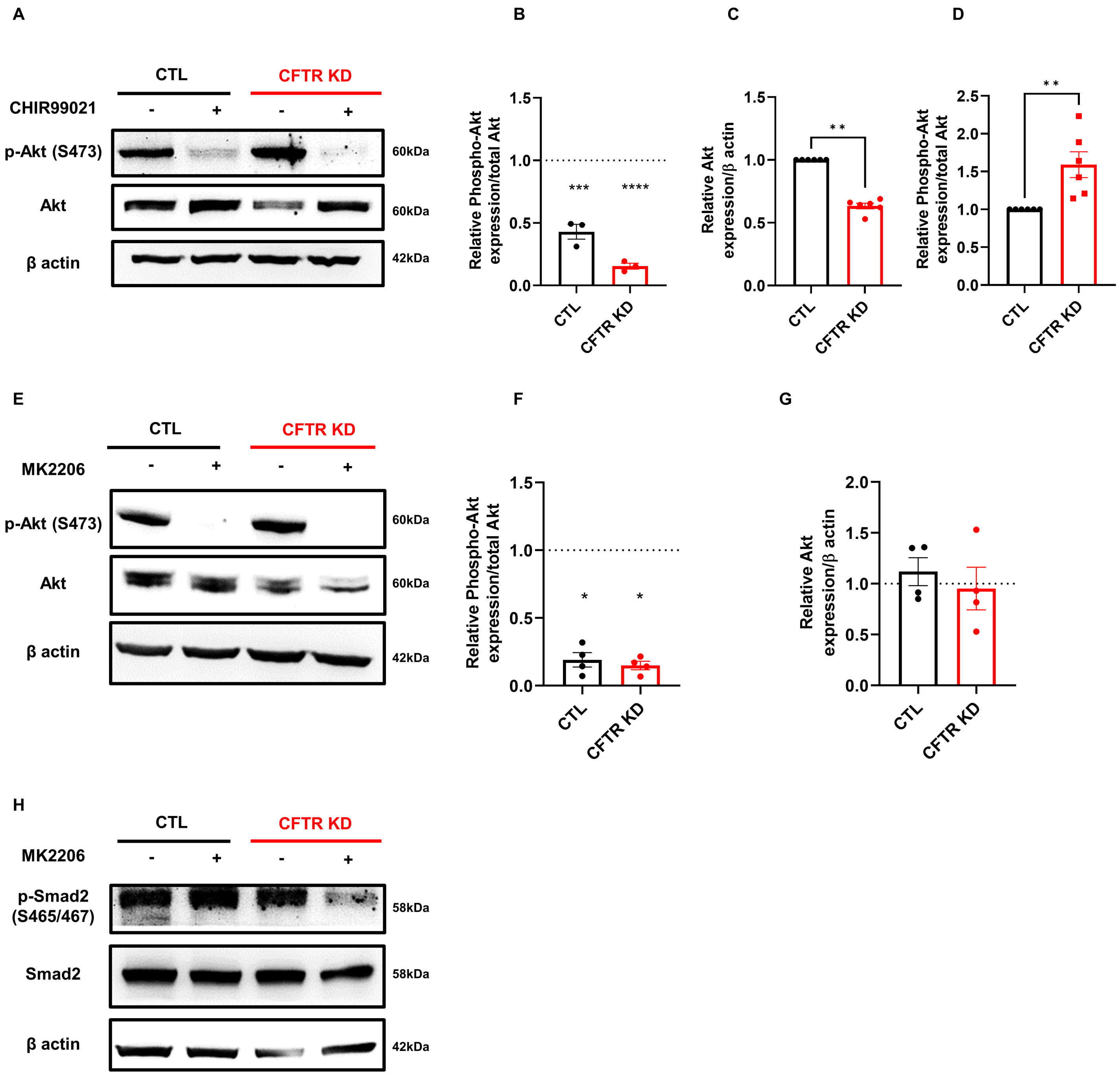

| | Age | Sex | Smoker | Pathology | Mutation | HAECs[1] | TEER[2] ($\Omega.cm^2$) | Cilia beating frequency (Hz) |
|---|---|---|---|---|---|---|---|---|
| CF MD048502 | 25 | Unknown | No | CF | Homozygous F508del | Bronchial | 411+/-10 | 7.2+/-0.2 |
| CF MD045303 | 35 | Male | No | CF | Homozygous F508del | Bronchial | 412+/-12 | 7.5+/-0.1 |
| CF MD044502 | 25 | Female | No | CF | Homozygous F508del | Bronchial | 419+/-10 | 7.4+/-0.2 |
| CF MD0567 | 39 | Female | No | CF | Homozygous F508del | Bronchial | 528+/-16 | 7.5+/-0.1 |
| CF MD0487 | 24 | Male | No | CF | Heterozygous F508del/2183AA>G | Bronchial | 324+/-20 | 8.6+/-0.1 |
| CF MD0408 | 24 | Unknown | No | CF | Homozygous F508del | Bronchial | 311+/-22 | 8.0+/-0.1 |
| CF MD0638 | 39 | Male | No | CF | Heterozygous F508del/2789+5G>A | Bronchial | 397+/-07 | 8.6+/-0.1 |
| CF MD0437 | 27 | Male | No | CF | Homozygous F508del | Bronchial | 456+/-07 | 7.1+/-0.1 |
| CF MD0607 | 21 | Female | No | CF | Homozygous F508del | Bronchial | 397+/-09 | 7.9+/-0.1 |
| NCF MD056001 | 66 | Male | No | No pathology reported | - | Bronchial | 400+/-11 | 7.2+/-0.1 |
| NCF MD053701 | 61 | Male | No | No pathology reported | - | Bronchial | 431+/-09 | 7.3+/-0.1 |
| NCF MD0459000 | 70 | Female | No | No pathology reported | - | Bronchial | 324+/-10 | 7.9+/-0.1 |
| NCF MD020102 | 67 | Female | No | No pathology reported | - | Bronchial | 356+/-09 | 7.1+/-0.2 |
| NCF MD0787 | 56 | Female | No | No pathology reported | - | Bronchial | 295+/-17 | 11.2+/-0.5 |
| NCF MD0835 | 35 | Male | No | No pathology reported | - | Bronchial | 210+/-08 | 7.9+/-0.6 |
| NCF MD0801 | 27 | Male | No | No pathology reported | - | Bronchial | 229+/-04 | 8.3+/-0.5 |
| NCF MD0720 | 41 | Male | No | No pathology reported | - | Bronchial | 206+/-06 | 8.5+/-0.6 |

| Gene | Forward (5’ to 3’) | Reverse (3’ to 5’) |
|---|---|---|
| TGF-β1 | CCCTGGACACCAACTATTGC | TGCGGAAGTCAATGTACAGC |
| DKK1 | AGCGTTGTTACTGTGGAGAAG | GTGTGAAGCCTAGAAGAATTACTG |
| β-catenin | TCGCCAGGATGATCCCAGC | GCCCATCCATGAGGTCCTG |
| Fibronectin | CACGGGAGCCTCGAAGAG | ACAACCGGGCTTGCTTTG |
| Smad2 | ATCCTAACAGAACTTCCGCC | CTCAGCAAAAACTTCCCCAC |
| 18S | GTAACCCGTTGAACCCCATT | CCATCCAATCGGTAGTAGCG |

| Reagents or Resource | Source | Identifier |
|---|---|---|
| **Experimental models** | | |
| NCF Primary HAECs (See supplementary table 1) | Epithelix Sàrl | EP01MD |
| CF Primary HAECs (See supplementary table 2) | Epithelix Sàrl | EP07MD |
| Calu-3 cell line | ATCC | HTB-55 |
| SBE-expressing cell line | Mouse Embryonic Fibroblasts (Gift from Prof. B. Wehrle-Haller) | PMID: 16549026 |
| **CRISPR products** | | |
| Cas9 Lentiviral particles | GeneCopoeia | LPP-CP-LVC9NU-10-100-cs |
| Scramble sgRNA Lentiviral particles | GeneCopoeia | LPPCCPCTR01L03-100-cs |
| **RNAscope® resources** | | |
| RNAscope® Multiplex Fluorescent Reagent Kit v2 | Advanced Cell Diagnostics | 323135 |
| RNAscope® Probe - Hs-TGFB1-C2 - Homo sapiens transforming growth factor beta 1 (TGFB1) mRNA | Advanced Cell Diagnostics | 400881-C2 |
| RNAscope® Probe - Homo sapiens dickkopf WNT signaling pathway inhibitor 1 (DKK1) mRNA | Advanced Cell Diagnostics | 421411-C3 |
| HybEZ oven | Advanced Cell Diagnostics | 321720 |
| Opal™ 520 Reagent Pack | Akoya Biosciences | FP1487001KT |
| Opal™ 570 Reagent Pack | Akoya Biosciences | FP1488001KT |
| Opal™ 690 Reagent Pack | Akoya Biosciences | FP1497001KT |
| **TGF-β bioassays** | | |
| Phospha-lightTM assay system | ThermoFisher | T1015 |
| **Pharmacological drugs** | | |
| Cycloheximide | Sigma | 01810 |
| CHIR99021 | Sigma | SML1046 |
| SB431542 | Sigma | 616461 |
| MK2206 | Seleckchem | S1078 |
| **Antibodies** | | |
| DKK1 | Cell Signaling | 4687S |
| Phospho-β-catenin (Ser552) | Cell Signaling | 9566S |
| β-catenin | Cell Signaling | 9562S |
| Fibronectin | Polyclonal antisera against human | Clone 1801 |

| | | |
|---|---|---|
| | plasma fibronectin (Gift from Prof. B. Wehrle-Haller) | |
| Total Smad2 | Cell Signaling | 5339S |
| Phospho-Smad2 (Ser465/467) | Cell Signaling | 3108S |
| Phospho-Smad2 (Ser465/467)/Smad3 (Ser423/425) | Cell Signaling | 8828S |
| Total Smad2/Smad3 | Cell Signaling | 3102S |
| GAPDH | Millipore | MAB374 |
| β-actin | Sigma | A1978 |
| Smad7 | Abcam | AB216428 |
| Total Akt | Cell Signaling | 9272S |
| Phospho-Akt (Ser473) | Cell Signaling | 9271S |
| Ki67 | DAKO | M7240 |
| Goat anti-Rabbit HRP | Sigma | A8275 |
| Goat anti-Mouse HRP | Sigma | A5278 |
| Alexa Fluor™ 568 goat anti-rabbit (H+L) | ThermoFisher | A11011 |
| Alexa Fluor™ 647 goat anti-rabbit (H+L) | ThermoFisher | A21245 |
| Alexa Fluor™ 568 goat anti-mouse (H+L) | ThermoFisher | A11031 |
| Alexa Fluor™ 647 goat anti-mouse (H+L) | ThermoFisher | A21236 |
| **Oligonucleotides** | | |
| qPCR primers | Microsynth | See supplementary table 2 |
| **Other reagents** | | |
| MEM-Glutamax | ThermoFisher | 41090-028 |
| MucilAir Culture Medium | Epithelix Sàrl | EP04MM |
| DMEM high glucose | Sigma | D6429 |
| Hygromycin B | Corning | 30-240-CR |
| Non-Essential Amino Acids | Bioconcept | 5-13K00-H |
| HEPES | ThermoFisher | 15630-056 |
| Sodium pyruvate 100X | ThermoFisher | 11360-039 |
| Fetal Bovine Serum (FBS) | ThermoFisher | 10270-106 |
| Penicillin / Streptomycin / Fungizone | Bioconcept | 4-02F00-H |
| BSA | Applichem | A1391 |
| Trypsin-EDTA 10X | ThermoFisher | 15400-054 |
| PBS | ThermoFisher | 14190-094 |
| Nonidet-P40 | Applichem | A1694 |

| | | |
|---|---|---|
| cOmplete Protease Inhibitor Cocktail | Roche | 04693124001 |
| Pierce BCA protein assay kit | ThermoFisher | 23228 |
| Porablot NCP nitrocellulose membrane | Macherey-Nagel | 741280 |
| Tween® 20 | Sigma | P2287 |
| SuperSignal™ West Pico PLUS Chemiluminescent Substrate | ThermoFisher | 34580 |
| Paraformaldehyde (PFA) | Sigma | 158127 |
| Triton 100X | Sigma | T8787 |
| DAPI | Applichem | A4099 |
| RNeasy mini kit | Qiagen | 74106 |
| PowerUp SYBR Green Master Mix | Appliedbiosystems | A2574 |
